# Endometrial Hyperplasia Risk Is Increased by High-Fat Diet Via Estrogen-Driven Stromal Fibroblast Reprogramming Toward a Pro-Fibrotic State

**DOI:** 10.64898/2026.03.20.713224

**Authors:** Hilary J. Skalski, Abigail Z. Bennett, Lauren E. Wood, Shannon K. Harkins, Amelia R. Arendt, Amanda G. Lopez Espinosa, Gregory W. Burns, Emmanuel N. Paul, Galen Hostetter, Katelyn Becker, Marc Wegener, Marie Adams, Jose M. Teixeira, Kin Lau, Ronald L. Chandler

## Abstract

The uterine endometrium is capable of scarless regeneration under coordinated estrogen and progesterone signaling across the menstrual cycle. Obesity suppresses progesterone production, leading to chronic estrogen exposure and increased endometrial hyperplasia (EH) risk. To define how obesity alters endometrial cell states, endometrial tissues from control and EH-predisposed mice fed either a control diet or a high-fat diet (HFD) were analyzed by single-cell RNA sequencing and tissue phenotyping. HFD reprogrammed endometrial stroma towards an inflammatory, pro-fibrotic state, reducing progesterone receptor-network-associated Aldh1a2^+^ fibroblasts and expanding estrogen receptor–network-associated Gsn⁺ fibroblasts. HFD further impaired macrophage recruitment and promoted hyperplastic epithelial signatures, consistent with increased disease severity in an EH mouse model. Stromal deletion of Estrogen Receptor α established stromal estrogen signaling as a driver of HFD-induced extracellular matrix (ECM) accumulation. Collectively, these findings identify HFD-driven fibroblast reprogramming as a central mechanism linking estrogen dominance to stromal fibrosis, defective immune clearance, and heightened EH susceptibility. We propose that, in response to progesterone, fibroblast-mediated ECM remodeling is vital to normal endometrial homeostasis.

**Graphical Abstract.**
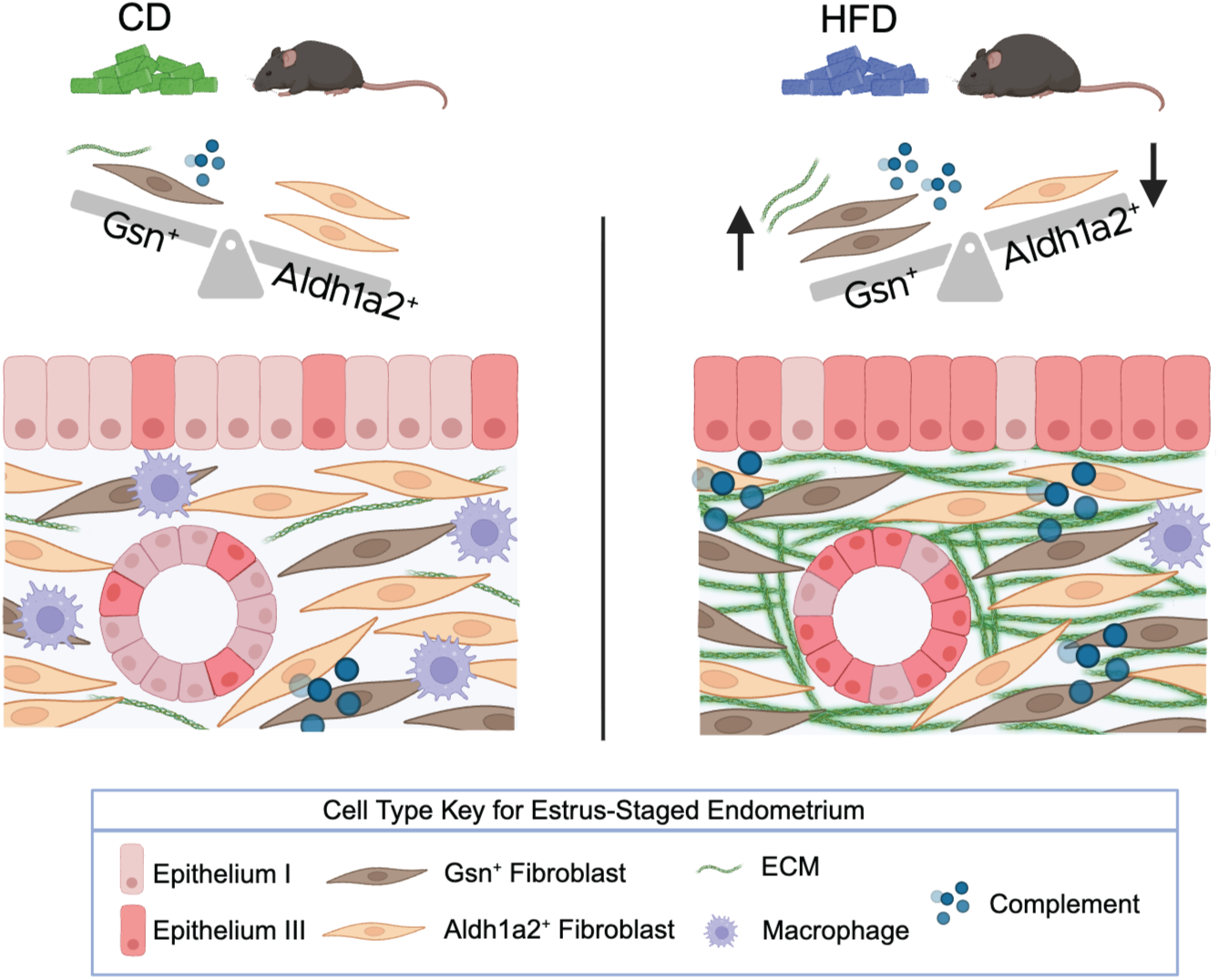
HFD-induced estrogen dominance disrupts endometrial fibroblast homeostasis to predispose the endometrium to disease. This study demonstrates that HFD drives estrogen-dependent reprogramming of stromal fibroblasts, characterized by inflammation, stromal ECM accumulation and fibrosis, and a post-ovulatory shift from PGR-network-associated Aldh1a2^+^ Fibroblasts toward increasing ER-network-associated Gsn^+^ Fibroblasts. These fibroblast changes are accompanied by a reduction in endometrial macrophages and a transcriptomic shift of HFD epithelium toward hyperplastic epithelium seen in a mouse model of EH. Figure made with BioRender.

## Introduction

In cycling females, the endometrium undergoes continual scarless regeneration and remodeling^1^. The bulk of the endometrium is composed of stromal fibroblasts and the extracellular matrix (ECM), which support the adjacent single columnar layer of luminal and glandular epithelium and are essential for successful embryo implantation and pregnancy maintenance. Cyclicity and endometrial cell composition and function are hormonally regulated by the ovarian steroid hormones estrogen and progesterone. In the first half of the cycle, the proliferative phase, estrogen induces epithelial and stromal fibroblast proliferation^1^. During the last half of the cycle, the secretory phase, postovulatory progesterone prepares the endometrium for potential embryo implantation. This involves an increase in natural killer (NK) cells to promote immune tolerance, along with epithelial glycogen production and release, stromal edema, and, in humans, subsequent fibroblast decidualization^1–4^. In the absence of embryo implantation, falling estrogen and progesterone levels trigger menses— the breakdown of the endometrium. This process, along with subsequent repair, involves coordinated immune system activity, including macrophage infiltration^1,5^.

Estrogen and progesterone signaling influence cellular crosstalk between endometrial stroma fibroblasts, epithelium and immune cells. While Estrogen Receptor α (ERα; *Esr1*) is expressed in both the endometrial epithelium and stroma, estradiol indirectly stimulates epithelial cell proliferation through fibroblast ERα signaling and subsequent fibroblast-derived paracrine factors^6–9^. To prevent excessive epithelial proliferation, another role of progesterone is to regulate estrogen’s proliferative effects^10,11^. Both the endometrial epithelium and fibroblasts express the progesterone receptor (PGR) and signaling through both cell types is vital to controlling estrogen-induced epithelial proliferation^1,9,12–14^. Uterine immune cells lack receptors for estrogen and progesterone, therefore it’s hypothesized that immune cells respond to estrogen and progesterone indirectly through epithelial and fibroblast cytokines^3,15^. Hormone imbalances can therefore lead to dysregulated cell-cell signaling, resulting in excessive endometrial epithelial proliferation and impairment of normal cyclical processes including endometrial remodeling, shedding, and repair^16,17^. As a result, the loss of normal endometrial cyclicity presents as abnormal menstrual bleeding and increases risk of endometrial diseases, including endometrial hyperplasia (EH) and endometrial cancer (EC)^16,18–20^.

In women with obesity, disruption of the central hypothalamic-pituitary-gonadal (HPG) axis contributes to abnormal ovarian steroid hormone production, anovulation, and menstrual dysfunction^21–23^. For example, altered levels of adipose-derived adipokines can disrupt hypothalamic kisspeptin neurons and subsequent gonadotropin-releasing hormone (GnRH) release and sensitivity^24,25^. Consequently, pituitary luteinizing hormone (LH) amplitude is decreased in women with obesity^26^. A burst of LH, or an LH surge, is needed to induce ovulation and generate the corpora lutea (CL), which produces progesterone^21–23^. Diminished LH therefore dampens progesterone secretion. Women with obesity have lower urinary levels of progesterone metabolite pregnanediol glucuronide (Pdg)^26^. They are also more likely to have longer menstrual cycles, often with a lengthened estrogen-dominant proliferative phase and shortened progesterone-dominant secretory phase^27^. Mice treated with a 60% high-fat diet (HFD) exhibited prolonged estrous cycles, characterized by an increased frequency in the estrogen-dominant first half of the cycle compared to the progesterone-dominant latter half^28^.

To understand the impact of obesity on endometrial cell types and its role in EH pathogenesis, we placed both control mice and mice genetically predisposed to develop EH on either a control diet (CD) or a HFD and performed endometrial single-cell RNA sequencing (scRNA-seq) and phenotypic analysis. We reveal that HFD disrupts endometrial homeostasis by reprogramming the stromal fibroblast transcriptome into inflammatory and pro-fibrotic states. Moreover, HFD reduced the amount of progesterone receptor (PGR)-network-associated Aldh1a2^+^ Fibroblasts, leading to a relative expansion of inflammatory estrogen receptor (ER)-network-associated Gsn^+^ Fibroblasts in the endometrium. Using the *Esr1^fl/fl^; Amhr2^Cre/+^* mouse model, we established that stromal ER signaling drives HFD-induced ECM accumulation. We further found that HFD reduced macrophage recruitment and dysregulated genes that suppress hyperplastic disease formation. This implies that HFD-induced fibroblast dysfunction induces persistent tissue damage by inhibiting immune cell clearance and predisposes the epithelium to hyperplastic disease. When using a genetically predisposed mouse model of EH, we showed that HFD-associated stromal fibrosis and reduced macrophages persist, and further increases EH disease severity. Therefore, we propose that a predominant inflammatory ER-network-associated Gsn^+^ fibroblast induces ECM accumulation with aberrant repair, thus predisposing women to endometrial disease, including EH.

## Results

### scRNA-seq of endometrial cells indicates HFD-induced fibroblast dysregulation

In women, obesity can disrupt the balance of estrogen to progesterone, increasing risk for abnormal menstrual bleeding, EH, and EC^16,18–20,26,27^. Consistent with observations in obese women, we previously found that mice fed a HFD exhibit reduced serum progesterone levels and lengthened estrous cycles, with an increased prevalence of the estrus stage^28^. However, these mice do not develop histologically evident EH^28,29^. Despite the absence of histological changes, HFD induces transcriptomic alterations in the endometrium^29^. Because *Pten* loss-of-function mutations—while present in normal endometrium—are more commonly associated with EH^30–33^, introducing this mutation provides a sensitized model to uncover how obesity may drive EH at the molecular level. Therefore, to investigate the effects of HFD on the endometrium in the context of EH development and progression, we performed scRNA-seq using an established EH mouse model^34^ with a targeted *Pten* loss-of-function mutation driven by *Pax8^Cre^*. *Pten floxed* control (*Pten^floxed^;Pax8^+/+^*), heterozygous (*Pten^fl/+^;Pax8^Cre/+^*), and homozygous (*Pten^fl/fl^;Pax8^Cre/+^*) mutant mice were placed on CD or HFD for a minimum of 21.5 weeks (Fig. 1a; Supplementary Fig. 1a-d). In accordance with our previous results, all HFD-treated groups were glucose intolerant and weighed significantly more than the CD-treated groups (Supplementary Fig. 1d-g)^28,29^. Uterine weight was significantly increased with *Pten* mutation, which agrees with previous findings utilizing *Pax8^Cre^*^34^ (Supplementary Fig. 1h,i).

**Fig. 1.**
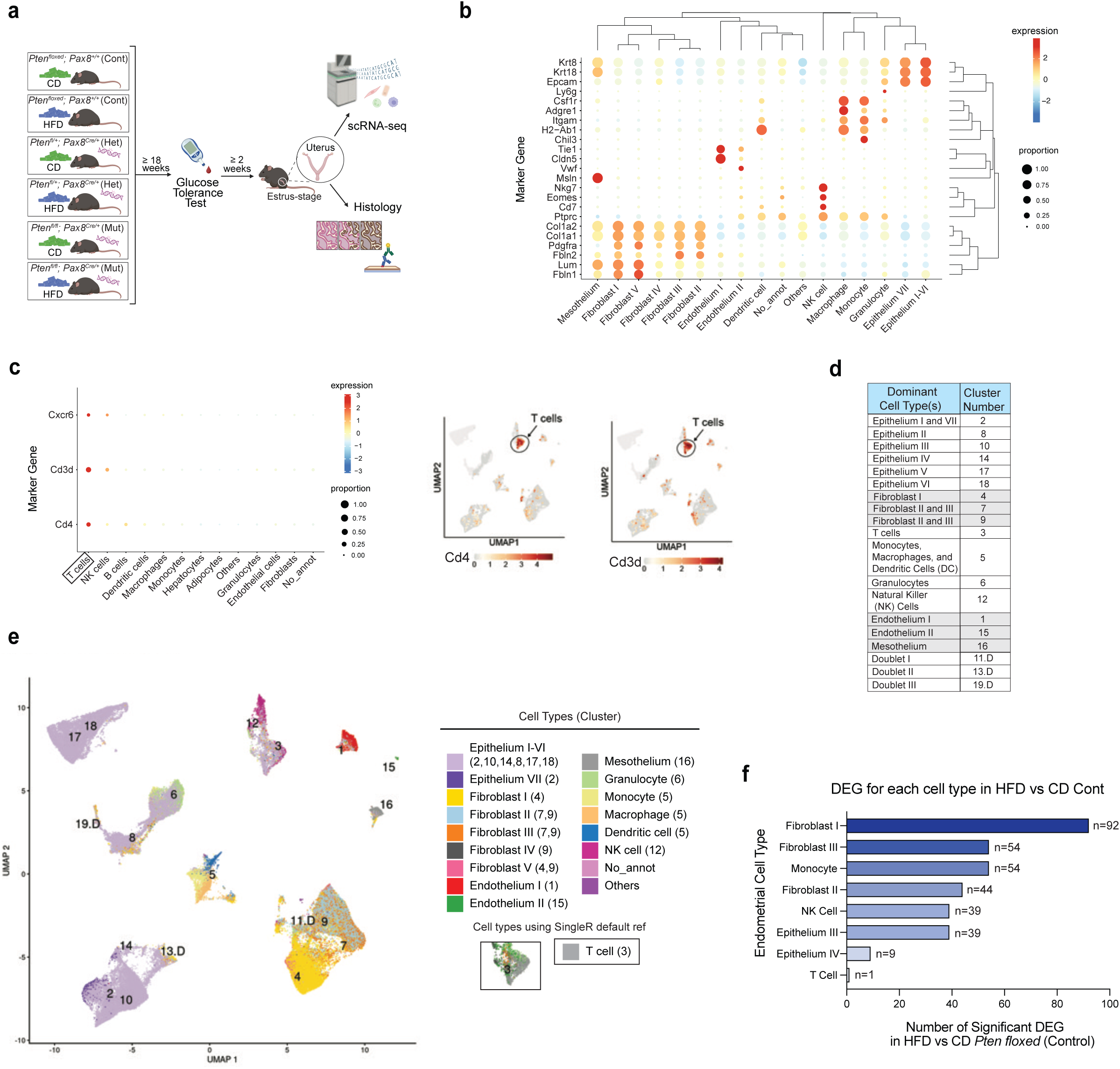
HFD leads to differential gene expression in several endometrial cell types, most predominantly in fibroblasts. (a) Schematic experimental timeline for scRNA-seq mouse cohort. Figure made with BioRender. (b,c) Bubbleplots of known marker genes for different identified cell types in CD and HFD controls using both the adult uterus dataset and the default reference for T cells. Additional UMAPs of T cell markers Cd3d and Cd4 showing that they are the most prominent in the default referenced T cells (Cluster 3). (d) Table of scRNA-seq identified clusters in the endometrium and the most commonly identified cell types found in each. (e) UMAP illustration of endometrial cell type clusters over all scRNA-seq conditions as predicted by SingleR with the mouse adult uterus dataset reference. T cells were predicted to be in Cluster 3 based on the default SingleR reference. Clusters 11.D, 13.D, and 19.D were identified as duplicates and were therefore removed from further analysis. (f) Number of DEGs in HFD versus CD *Pten^floxed^* (control) mice per endometrial cell type. DEG significance determined at FDR<0.05.

To minimize transcriptomic variability driven by hormonal fluctuations throughout the estrous cycle, all samples for scRNA-seq were collected specifically during the estrus stage from mice cycling asynchronously. To allow for isolations to be performed on different days, uteri were dissociated following our published tissue digestion method^29^ and live cells were fixed and processed using the 10x Genomics Chromium Single Cell Gene Expression Flex (v1) protocol. Using 10x Flex, an average of 4,905 reads per cell were sequenced with 1,800 genes detected per cell. Cell type annotation using SingleR with the mouse adult uterus dataset^35^ revealed 19 distinct clusters were present in the CD- and HFD-treated groups (Fig. 1b-e). Clusters 11.D, 13.D and 19.D were identified as containing a high proportion of doublets, which is denoted by a “D”, and therefore were subsequently excluded from further analysis (Supplementary Fig. 2a-c). We confirmed that identified cell types expressed published canonical markers (Fig. 1b,c)^36–40^. Epithelial cells (identified in Clusters 2, 10, 8, 14, 17, and 18) highly expressed *Krt8*, *Krt18*, and *Epcam* and were therefore labeled as Epithelium I-VII. Fibroblasts (identified in Clusters 4, 7, and 9) highly expressed *Fbln1*, *Fbln2*, *Lum*, *Pdgfra*, *Col1a1*, and *Col1a2* and thus labelled Fibroblasts I-V. Endothelial cells (identified in Clusters 1 and 15) highly expressed *Tie1*, *Cldn5*, and *Vwf* and were named Endothelium I and II. Mesothelium (identified in Cluster 16) highly expressed *Msln*. All immune cells expressed *Ptprc* (CD45) and myeloid cells expressed *Itgam* (CD11b). Natural Killer (NK) cells (identified in Cluster 12) highly expressed *Nkg7*, *Eomes*, and *Cd7*. Monocytes (identified in Cluster 5) highly expressed *Chil3*. Dendritic cells (DCs) (identified in Cluster 5) highly expressed *H2-Ab1* and *H2-Aa*. Macrophages (identified in Cluster 5) highly expressed *Adgre1* (F4/80) and *Csf1r*. Granulocytes (identified in Cluster 6) highly expressed *Ly6g*. Additional cell type annotation using the SingleR default reference dataset determined that Cluster 3 should be labelled as T cells (Fig. 1c-e). These T cells highly expressed T cell markers including *Cd4*, *Cd3d*, and *Cxcr6*. Epithelium V and VI (Clusters 17 and 18) were uniquely seen with *Pten* mutation and therefore, these clusters were identified as hyperplastic epithelial cells (Supplementary Fig. 3a,b). Additionally, the Cluster 6 cell population (granulocytes) was increased with *Pten* mutation, which agrees with previous findings in mouse models with targeted uterine *Pten* mutation^41^. When examining the transcriptomic effect of HFD alone (comparing HFD- to CD-treated controls), the highest number of DEGs were present in fibroblasts (Fig. 1f). DEGs from comparisons of the effect of diet in *Pten* heterozygous and homozygous mutant mice, as well as the effect of *Pten* mutation in CD and HFD conditions, were also examined (Supplementary Fig. 3c-h).

Due to the number of DEGs observed in comparisons between HFD and CD control fibroblasts, we further profiled all the three prominent fibroblast subtypes (Fibroblast I, II, and III). Marker genes were identified to more effectively distinguish between fibroblast clusters (Supplementary Fig. 4a). *Adamts1*, *Clec3b*, *Col3a1*, *Col14a1*, *Cxcl12*, *Dpt, Efemp1, Gsn*, *Has1*, *Lum*, *Mgp*, *Sparc*, and *Sparcl1* were among the top 50 marker genes for Fibroblast I (the dominant cell type in Cluster 4). Marker genes *Adamts1*, *Adh1*, *Col1a2*, *Col3a1*, *Clec3b*, *Col14a1, Dpt, Efemp1, Gpx3, Gsn*, *Has1*, *Lum*, *Mgp, Sparc,* and *Sprarcl1* are markers of mouse endometrial myofibroblasts derived from regenerative fibroblasts^36^, though *Acta2* (α-smooth muscle actin), a characteristic marker of myofibroblasts^42,43^, was not enriched in the Fibroblast I cell type. Fibroblast I additionally shares a significant amount of marker genes (n=23/50; 46%) with human endometrial “C7 fibroblasts”, including *Cxcl12*, *Dcn*, *Dpt*, *Gsn*, *Lum*, *Mgp*, *Serping1*, and *Timp2* (Supplementary Fig. 4b)^44^. Fibroblasts II and III alternatively comprise the majority of cells in both Clusters 7 and 9, which is consistent with their sharing more than half (n=27) of their top 50 marker genes— a set that is distinct from Fibroblast I. Fibroblasts II/III marker genes included *A2m*, *Aldh1a2*, *Ccl11*, *Col6a4*, *Dio2*, *Hsd11b2*, *Htra1*, and *Smoc2* (Supplementary Fig. 4a). These marker genes are enriched in mouse endometrial maturational fibroblasts associated with the second half of the estrous cycle, when progesterone is elevated^36^. From the top marker genes, Fibroblast I (Cluster 4) will be referred to as “Gsn^+^ Fibroblast”, whereas Fibroblast II/III (Clusters 7/9) will be referred to as “Aldh1a2^+^ Fibroblast”, due to a combination of high gene expression levels and a high proportion of cells expressing *Gsn* and *Aldh1a2*, respectively (Supplementary Fig. 4c). The comprehensive top 200 genes changing across the three fibroblast clusters were also determined (Supplementary Fig. 4d). Transcription factor (TF) activity analysis was additionally performed on the three fibroblast clusters (Supplementary Fig. 4e). In both CD and HFD, the ER network (TFs ESR1 and ESR2) is predicted to be active in the Gsn^+^ Fibroblast. In contrast, the PGR network is predicted to be an enriched target in the Aldh1a2^+^ Fibroblast, consistent with marker genes that indicate the Aldh1a2^+^ cells are enriched during progesterone-dominant stages of the cycle.

### HFD fibroblast enrichment of complement, ECM, mitochondrial dysfunction, and cell-matrix adhesion

We then examined how the three stromal fibroblast populations changed with diet. HFD led to a decrease in PGR-network-associated Aldh1a2^+^ Fibroblasts, thereby resulting in a reduced proportion of these cells relative to the ER-network-associated Gsn^+^ Fibroblast (Fig. 2a). We proceeded to analyze gene expression changes induced by HFD in these fibroblast subtypes. The majority (n=111; 75%) of all 148 DEGs in these clusters were upregulated (Fig. 2b,c). The Gsn^+^ Fibroblast had 60 upregulated and 14 downregulated DEGs that were unique to this cell type, while Aldh1a2^+^ Fibroblast II and III share a considerable number (n=22/51; ∼43%) of upregulated DEGs. Further analysis of the Gsn^+^ Fibroblast, which accounted for 92 of the 148 fibroblast DEGs, similarly demonstrated that the majority (n=76; ∼83%) of DEGs were upregulated with HFD (Fig. 2d). Upregulated genes included several components of the classical complement pathway including *C1ra*, *C1s1*, *C3*, *C4b*, and *C7*. Tight regulation of the complement system is essential, as excessive activation can result in chronic inflammation and subsequent tissue damage^45,46^. HFD also upregulated *Il6,* a proinflammatory cytokine upregulated in fibrosis induced by chronic inflammation^47^. Additional immune-related genes included downregulation of *Il15*, an cytokine released from decidualized fibroblasts to induce uterine NK (uNK) cell maturation^48^, and an upregulation of the *Cd47* “don’t eat me” gene^49^. Pathologically, *Cd47* prevents macrophage phagocytic activity and is upregulated in fibroblasts with lung fibrosis^49^, EH, and EC^50^. Moreover, extracellular matrix and tissue remodeling genes *Clec3b*, *Dcn*, *Ecm1*, *Ecm2*, *Efemp1, Fbln1*, *Il6, Sod3, Sned1,* and *Vwa1* were upregulated with HFD. In the postovulatory portion of the menstrual cycle, progesterone induces endometrial ECM remodeling to prepare the tissue for embryo implantation^51–53^. Gene ontology (GO) enrichment analysis of Gsn^+^ Fibroblast DEGs likewise displayed GO terms involving the immune response and extracellular matrix (Fig. 2e,f). In the Aldh1a2^+^ Fibroblast II, HFD led to 44 DEGs including an upregulation of *Fgf10* (Fig. 2g). Fibroblast growth factors (FGFs) are known mitogens for epithelial cells^54^. GO enrichment analysis from Aldh1a2^+^ Fibroblast II DEGs showed the highest enrichment for mitochondrial depolarization (Fig. 2h). Obesity induces oxidative stress within endometrial stromal cells, leading to mitochondrial dysfunction^55^. The Aldh1a2^+^ Fibroblast III population had 54 DEGs (Fig. 2i), including *Clec3b* upregulation, which was also observed in the Gsn^+^ Fibroblast. Enrichment analysis of the Aldh1a2^+^ Fibroblast III displayed GO terms involving cell-matrix adhesion, including a downregulation of *Pcsk5* (Fig. 2j). Cell-matrix adhesion is not only vital to tissue organization, but also to the regulation of immune responses and wound healing^56^. Comparable to Gsn^+^ Fibroblast, Aldh1a2^+^ Fibroblasts II and III also upregulated extracellular matrix genes *Dcn, Ecm2, Sned1,* and *Adamtsl1* (Fig. 2d,g,i). *Sod3* was significantly upregulated in all three fibroblast populations with HFD (Fig. 2d,g,i). Superoxide dismutase (*Sod3*) is an extracellular antioxidant enzyme upregulated in adipose tissue of HFD-treated mice and is thought to protect against HFD-induced oxidative damage^57^. Moreover, while *Gsn* is a marker for the Gsn^+^ Fibroblast I, HFD upregulated *Gsn* expression in all three fibroblast subtypes (Fig. 2d,g,i). Gelsolin is associated with pulmonary inflammation-associated fibrotic disease^58^. These enrichment analyses align with the pathological outcome that chronic inflammation drives altered stromal homeostasis, resulting in ECM dysfunction^59^.

**Fig. 2.**
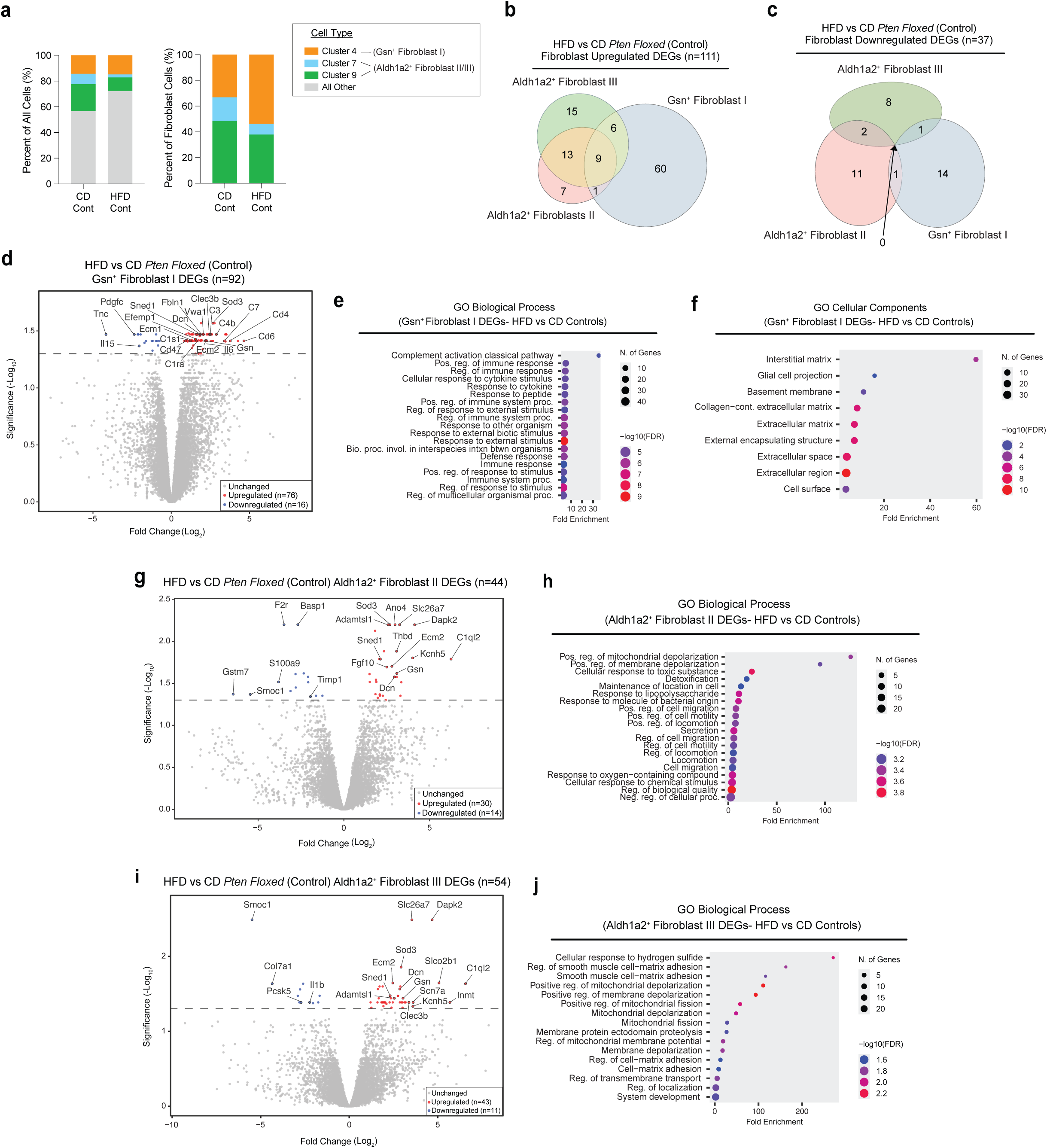
HFD-induced fibroblast imbalance and dysregulation of ECM, inflammation, mitochondrial depolarization, and cell-matrix adhesion. (a) Stacked bar graphs comparing the CD and HFD proportions of the three major fibroblast subtypes (the Gsn^+^ Fibroblast and Aldh1a2^+^ Fibroblasts II and III) in *Pten^floxed^* (control) mice. (b,c) Euler diagram displaying the numbers of HFD-associated upregulated (n=111) or downregulated (n=37) DEGs present in the three fibroblast subtypes and if they are cell subtype specific or shared in *Pten^floxed^*(control) mice. (d) Volcano plot of HFD versus CD Gsn^+^ Fibroblast DEGs (n=92). Significantly upregulated genes (n=76) are in red and downregulated (n=16) are in blue. (e,f) Biological Processes or Cellular Components GO enrichment analysis dotplots from Gsn^+^ Fibroblast DEGs. (g) Volcano plot of HFD versus CD Aldh1a2^+^ Fibroblast II DEGs (n=44). Significantly upregulated (n=30) genes are in red and downregulated (n=14) are in blue. (h) Biological Processes GO enrichment analysis dotplots from Aldh1a2^+^ Fibroblast II DEGs. (i) Volcano plot of HFD versus CD Aldh1a2^+^ Fibroblast III DEGs (n=54). Significantly upregulated (n=43) genes are in red and downregulated (n=11) are in blue. (j) Biological Processes GO enrichment analysis dotplots from Aldh1a2^+^ Fibroblast III DEGs. DEG significance determined at FDR<0.05. Euler diagrams were made online with *eulerr.* Volcano plots were created online with VolcaNoseR^125^. GO enrichment plots were developed online with ShinyGO 0.82^126^.

In addition to DEGs, HFD also altered predicted TF activity in fibroblasts. HFD increased *Foxf2* and *Stat3* TF activity in both Gsn^+^ and Aldh1a2^+^ Fibroblasts (Clusters 4/9 and Clusters 4/7, respectively) and *Srf* in the Aldh1a2^+^ Fibroblast (Clusters 7 and 9) (Supplementary Fig. 4e). These proteins are involved in the development of tissue fibrosis^60–62^. Additionally, HFD decreased activity of *Klf4*, *Smad2*, and *Smad3*, and increased activity of *Klf17*. While histologically-evident EH is absent in HFD-treated control uteri, EH and EC are associated with the respective loss or gain of these proteins^63,64^. Disruption of the SMAD2/3 pathway is associated with fibroblast *Fgf10* upregulation (upregulated in Aldh1a2^+^ Fibroblast II) and enhanced epithelial cell proliferation and hyperplasia^65^.

### Increased endometrial stroma ECM and collagen deposition in response to HFD, indicating tissue fibrosis

Upregulation of inflammatory and ECM associated genes in stromal fibroblasts suggest HFD-induced endometrial inflammation and connective tissue accumulation. Chronic inflammation can drive fibroblast-mediated ECM deposition and fibrosis^59^. Histological sections of estrus-staged uteri revealed that HFD induced a marked increase in stromal ECM deposition (Fig. 3a-d), a key indicator of tissue fibrosis^66^. Additionally, HFD tissue showed elevated collagen—the primary ECM protein associated with fibrosis—in the endometrial stroma, further corroborating our findings of ECM accumulation and fibrotic changes (Fig. 3e-h). Moreover, given that gelsolin is associated with pulmonary inflammation-associated fibrotic disease^58^, is upregulated with HFD, and is a top marker of our Gsn^+^ (Gelsolin) Fibroblasts, we performed gelsolin staining on both CD and HFD endometrial samples. GSN^+^ fibroblasts were found throughout the CD- and HFD-treated endometrium, but was more prominent in the HFD-treated mice (Fig. 3i,j). Together, the increases in ECM, collagen, and gelsolin reinforce our scRNA-seq findings, supporting the interpretation that HFD-induced fibroblast dysregulation disrupts ECM homeostasis and drives fibrosis in the endometrium.

**Fig. 3.**
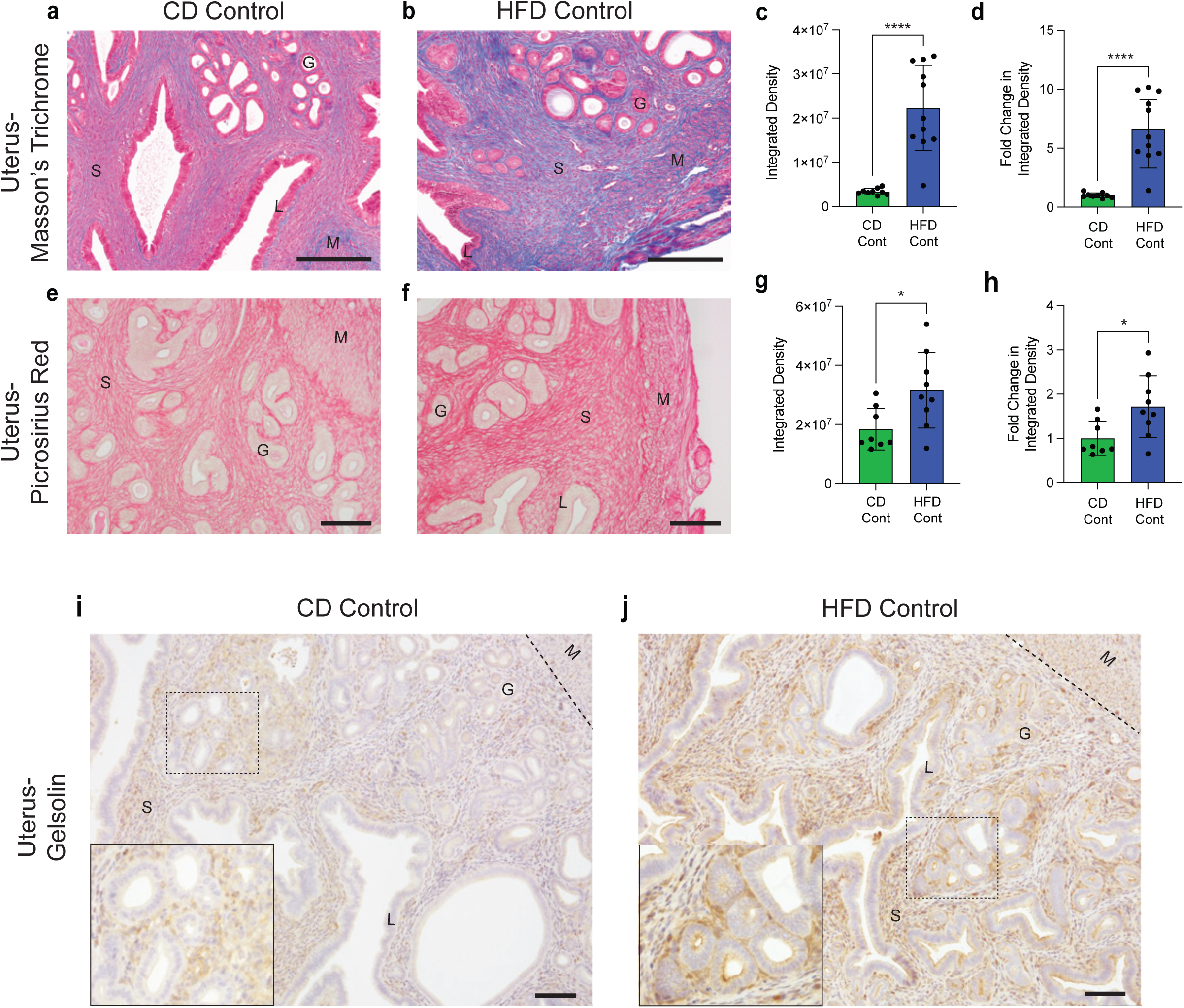
HFD leads to increased endometrial stroma ECM and collagen deposition, indicative of fibrosis. (a,b) Representative Masson’s Trichrome stain of mouse estrus-staged endometrial tissue for each diet. Blue color indicates ECM. Image= 20x magnification; scalebar=200 μm. (c,d) Quantification of CD and HFD trichrome slides using Image J with the Color Deconvolution plug-in. Graphs include raw integrated density of slides for each diet, as well as each CD- and HFD-induced fold change in integrated density compared to the CD average. Datapoints (CD n=9; HFD n=11) represent the average of four quantified sections from each mouse uterus (bar=mean ± s.d.). Two-tailed unpaired *t*-test statistic with Welch’s correction. (e,f) Representative picrosirius red stain for collagen in mouse estrus-staged endometrial tissue for each diet. Image= 20x magnification; scalebar=100 μm. (g,h) Quantification of CD (n=8) and HFD (n=9) picrosirius red-stained uteri was done as above for the trichrome-stained uteri. Two-tailed unpaired *t*-test statistic. (i,j) Representative Gelsolin IHC stain of estrus-staged endometrial tissue for each diet (CD n=8; HFD n=9). Main image= 10x magnification; inset=20x magnification; scalebar=100µm. M= myometrium; S= stroma; L= luminal epithelium; G=endometrial gland. Dashed line delineates the border between the endometrium and myometrium. *\*p<0.05; **p<0.01; ***p<0.001; ****p<0.0001*

### Disrupted stromal ERα signaling reduces HFD-associated ECM accumulation phenotype

Imbalances in estrogen and progesterone can disrupt normal cyclical processes of endometrial remodeling and repair^16,17^. HFD induces an unopposed estrogen state by suppressing progesterone production^28^, and obesity-associated estrogen dominance is the proposed mechanism by which obesity increases risk of endometrial diseases, including EH and EC^16,18–20^. To validate that HFD-associated increases in stroma collagen deposition were due to estrogen signaling in the fibroblasts, we utilized a conditional mouse model with stromal loss of ERα (*Esr1*)^67,68^. CD and HFD-treated *Esr1 floxed* control (*Esr1^floxed^;Amhr2^+/+^*) and homozygous mutant (*Esr1^fl/fl^;Amhr2^Cre/+^*) mice were placed on CD or HFD for a minimum of 21.5 weeks (Fig. 4a; Supplementary Fig. 5a-c). Consistent with our previous collections, CD- and HFD-treated control mice were collected in the estrus stage. On the other hand, in agreement with previous literature on ERα KO mice^69,70^, CD- and HFD-treated conditional *Esr1* mutants were collected during diestrus due to abnormal cycling (*data not shown*). In accordance with our previous HFD results, both HFD-treated groups were glucose intolerant and weighed significantly more than the CD-treated groups at the time of tissue harvest (Fig. 4b,c). As previously reported with reproductive tract targeted *Esr1* mutation, uteri of mutant animals were thin, and ovaries were enlarged with cystic hemorrhagic follicles and tumors (*not shown*)^71,72^. Endometrial ERα IHC-staining confirmed a decrease in ERα expression in stromal fibroblasts (Fig. 4d). Consistent with previous observations, histological samples from HFD control animals exhibited increased ECM deposition (Fig. 4e). With the additional *Esr1* mutation, HFD uteri exhibited ECM accumulation similar to that of the CD mutants, which was significantly less than the HFD control uteri. This suggests that estrogen signaling in endometrial fibroblasts is required for inducing the pathological ECM accumulation, and corresponding fibroblast dysfunction, seen with HFD.

**Fig. 4.**
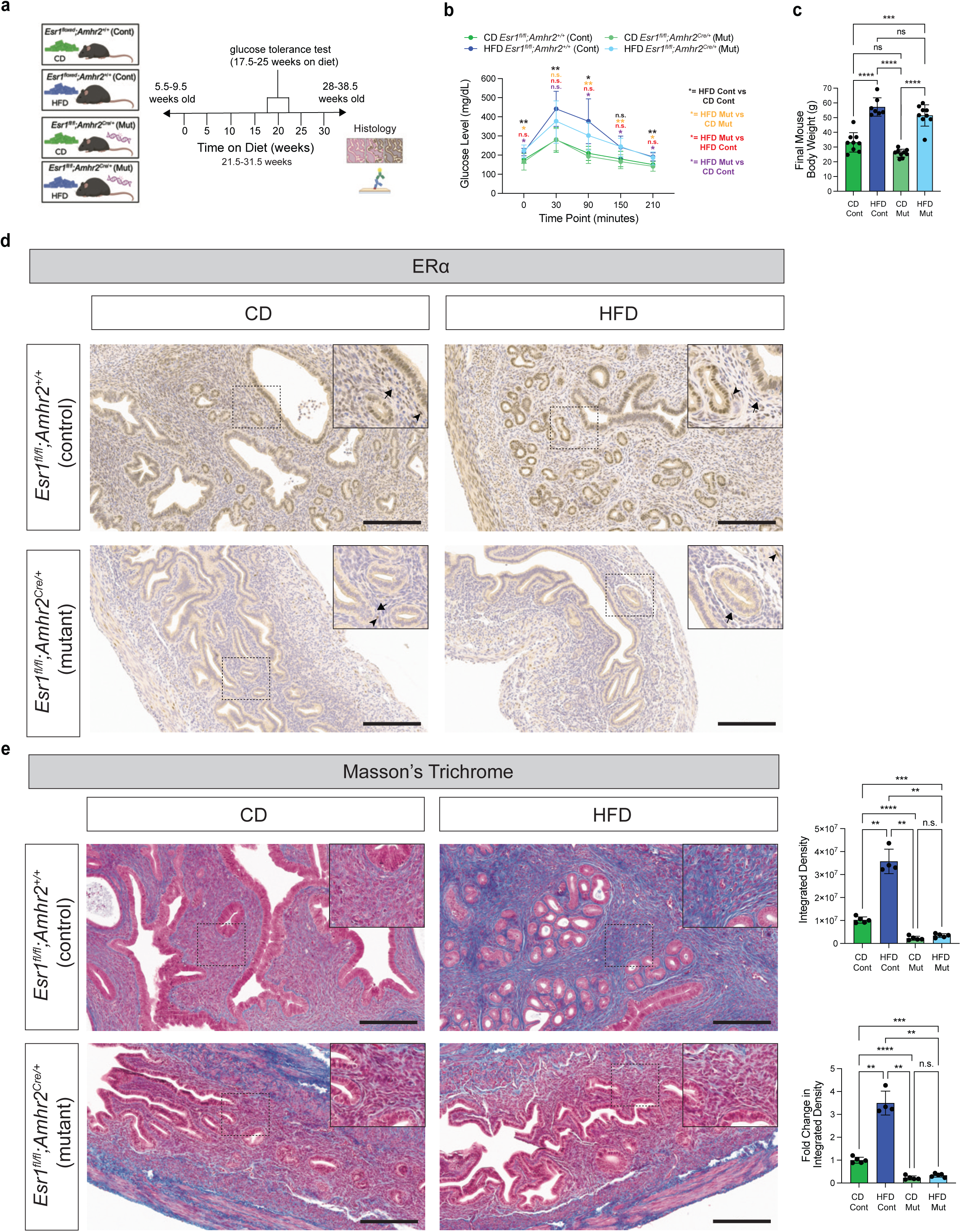
Disruption of endometrial fibroblast ERα signaling prevents HFD-induced ECM accumulation in the endometrium. (a) Experimental timeline for CD and HFD-treated *Esr1^floxed^; Amhr2^+/+^* (control) and *Esr1^fl/fl^; Amhr2^Cre/+^* (mutant) mice used for histology. Graphic made with BioRender. (b) Glucose tolerance test after 17.5-25 weeks on diet. Time point 0 fasting blood glucose (mg/dL) and post-glucose injection blood glucose levels were measured (30, 90, 150, and 210 minutes). CD (n=9) and HFD (n=8) control, and CD (n=9) and HFD (n=9) mutant mice. Datapoints represent a diet’s mean ± s.d. Two-way ANOVA with Geisser-Greenhouse correction and Tukey multiple comparison post hoc correction test statistic. (c) Final mouse body weight (g) of *Esr1^fl/fl^; Amhr2^+/+^* (cont) and *Esr1^fl/fl^; Amhr2^Cre/+^* (mut) mice on both CD and HFD. Measurement (datapoint) represents each mouse (bar=mean ± s.d.). Brown-Forsythe and Welch ANOVA with Dunnett T3 multiple comparison test statistic. (d) Representative ERα IHC stain of endometrial tissue from *Esr1* homozygous controls (CD=5; HFD=4) and mutants (CD=5; HFD=5) for each diet. Main image= 20x magnification; inset=40x magnification; scalebar=200 μm. Arrowhead indicates positive fibroblast staining and arrow indicates negative fibroblast staining. (e) Representative Masson’s Trichrome stain of endometrial ECM from *Esr1* homozygous controls (CD=5; HFD=4) and mutants (CD=5; HFD=5) mutants for each diet. Main image= 20x magnification; inset=40x magnification; scalebar=200 μm. Accompanying graphs quantifying ECM raw integrated density or fold change in integrated density in *Esr1* homozygous control and mutant endometrial stroma using Image J with the Color Deconvolution plugin. Samples were from CD- (n=5) or HFD- (n=4) treated control animals and CD- (n=5) or HFD- (n=5) treated mutant animals. Measurement (datapoint) represents each mouse’s representative sample mean (bar=mean ± s.d.). Brown-Forsythe and Welch ANOVA with Dunnett T3 multiple comparison test statistic. Tissues in histological samples were collected in the estrus stage and diestrus stage for CD- and HFD-treated controls, and CD- and HFD-treated *Esr1* conditional knockouts (cKO), respectively. *\*p<0.05; **p<0.01; ***p<0.001; ****p<0.0001*

### HFD leads to decreased NK cell activity and downregulation of monocyte migration-related gene expression, likely leading to decreased endometrial macrophages

Fibroblasts are central regulators of local immunity, driving conditions including chronic inflammation and cancer-associated immunosuppression^73^. The enrichment of HFD-induced immune-associated DEGs in fibroblasts prompted us to examine immune cells. After fibroblasts, the cell types with the highest number of HFD-induced DEGs were monocytes and natural killer cells (NK) (Fig. 1f). When examining the 54 monocyte DEGs, the majority (70.4%) were downregulated including monocyte recruitment and differentiation genes *Spp1*, *Mmp14,* and *Cxcl16* (Fig. 5a). GO Biological Processes enrichment analysis similarly presented DEGs related to cell migration and motility (Fig. 5b). Monocytes circulate in the blood and enter tissues where they then differentiate into macrophages or dendritic cells (DCs)^74^. Macrophages are prevalent during endometrial breakdown and repair and are known to remove endometrial debris, regulate inflammation, and promote endometrial repair^5,75^. Therefore, due to the downregulation of the monocyte-to-macrophage differentiation and chemoattractant genes, we examined the scRNA-seq data for changes in the proportion of macrophages with HFD. After subclustering Cluster 5, which contained a combination of monocytes, macrophages, and dendritic cells (Fig. 1d,e), we identified macrophages in subclusters 5.2 and 5.5 (Supplementary Fig. 6a,b). HFD-treated samples had a decrease in macrophages compared to CD-treated (Fig. 5c,d), which agrees with both the HFD-altered monocyte gene expression and dysregulated fibroblasts, which together may impede immune cell infiltration.

**Fig. 5.**
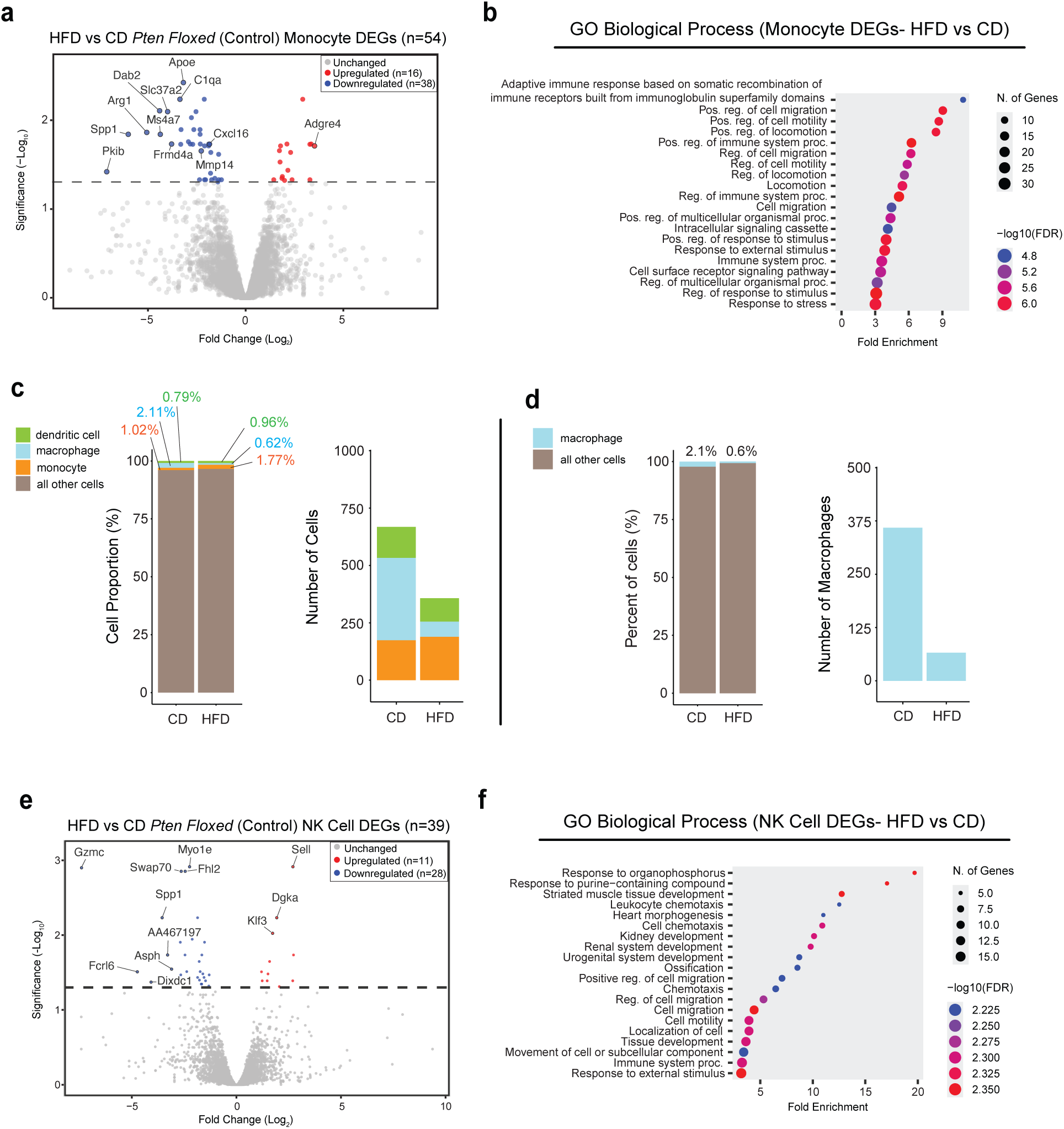
Monocyte and NK cell dysregulation and reduced uterine macrophages with HFD. (a) Volcano plot of HFD versus CD *Pten^floxed^* (control) mouse DEGs (n=54) in monocytes. Significantly upregulated (n=16) genes are in red and downregulated (n=38) are in blue. (b) Biological Processes GO enrichment analysis dotplot from monocyte DEGs. (c,d) Stacked bar graphs comparing either the proportions or total cell count of macrophages (subclusters 5.2 and 5.5) out of the total cell population in both CD and HFD. (e) Volcano plot of HFD versus CD DEGs in NK cells (n=39). Significantly upregulated (n=11) genes are in red and downregulated (n=28) are in blue. (f) Biological Processes GO enrichment analysis dotplot from NK cell DEGs. DEG significance determined at FDR<0.05. Volcano plots were created online with VolcaNoseR^125^. GO enrichment plots were developed online with ShinyGO 0.82^126^.

After monocytes, the immune cell type with the highest number of DEGs was NK cells (Fig. 1f). Uterine NK (uNK) cells are more commonly studied in pregnancy, where their functions include establishing immune tolerance^76^ and inducing vascular remodeling^77^. In the non-gravid endometrium, human uNK cells also regulate endometrial bleeding through structural changes to the spiral artery^52^. The majority (n=28/39; 71.8%) of NK cell DEGs were downregulated, with the top downregulated DEG being *Gzmc* (Granzyme C) (Fig. 5e). Downregulation of *Gzmc* suggests that HFD NK cells are in a resting state, as Granzyme C increases upon activation with IL-15^78^. *Il15* expression increases in endometrial stroma with rising progesterone levels and plays a critical role in uNK cell development, maturation, and proliferation^53^. Downregulation of *Il15* was observed in the HFD-treated Gsn^+^ Fibroblast population (Fig. 2d), suggesting that Gsn^+^ Fibroblasts may normally activate NK cells via IL-15 secretion. *Spp1*, in addition to its downregulation in monocytes, was also downregulated in NK cells, where its loss leads to impaired NK cell maturation and decreased cytolytic activity and cytokine secretion^79^. Enrichment analysis also revealed NK cell DEGs related to chemotaxis and cell migration (Fig. 5f). The final immune cell type with HFD-induced DEGs were T cells, where there was only 1 DEG (Fig. 1f)—an upregulation of *Sell*—which is upregulated in naïve T cells as compared to activated T cells.

### Epithelia from HFD-treated mice display a shift from stem cell-like signatures towards hyperplastic cell signatures seen in a conditional PTEN-deficient mouse model of EH

Because obesity is a risk factor for EH development^103^, we next wanted to examine the scRNA-sequencing data to see if the HFD control (*Pten^floxed^;Pax8^+/+^*) endometrial epithelial transcriptome more closely resembled the hyperplastic epithelium of our conditional PTEN-deficient mouse model of EH (*Pten^floxed^;Pax8^Cre/+^*). There were six epithelial clusters (Clusters 2, 10, 14, 8, 17, and 18) identified in our scRNA-sequencing data (Fig. 1d,e), which contained Epithelium I-VII. While Epithelium I composed the majority of Cluster 2 (Supplementary Fig. 7a), Epithelium VII was most commonly seen in that cluster (Supplementary Fig. 7b). Moreover, Epithelium V and VI (Clusters 17 and 18) only present in our PTEN-deficient mice (*Pten^floxed^;Pax8^Cre/+^*) (Supplementary Fig. 3a,b; Fig. 6a). Epithelium I was the most prominent epithelial cell type in CD control samples (*Pten^floxed^;Pax8^+/+^*), while Epithelium III dominated the HFD control samples (*Pten^floxed^;Pax8^+/+^*) (Fig. 6a). This HFD-associated prevalence of Epithelium III over Epithelium I persisted with heterozygous PTEN loss (*Pten^fl/+^;Pax8^Cre/+^*). Epithelium II increased in both CD and HFD PTEN heterozygous (*Pten^fl/+^;Pax8^Cre/+^*) and homozygous (*Pten^fl/fl^;Pax8^Cre/+^*) mice compared to the controls (*Pten^floxed^;Pax8^+/+^*). With homozygous PTEN loss (*Pten^fl/fl^;Pax8^Cre/+^*), hyperplastic Epithelium V and VI were the dominate epithelial subtypes in both CD and HFD samples.

**Fig. 6.**
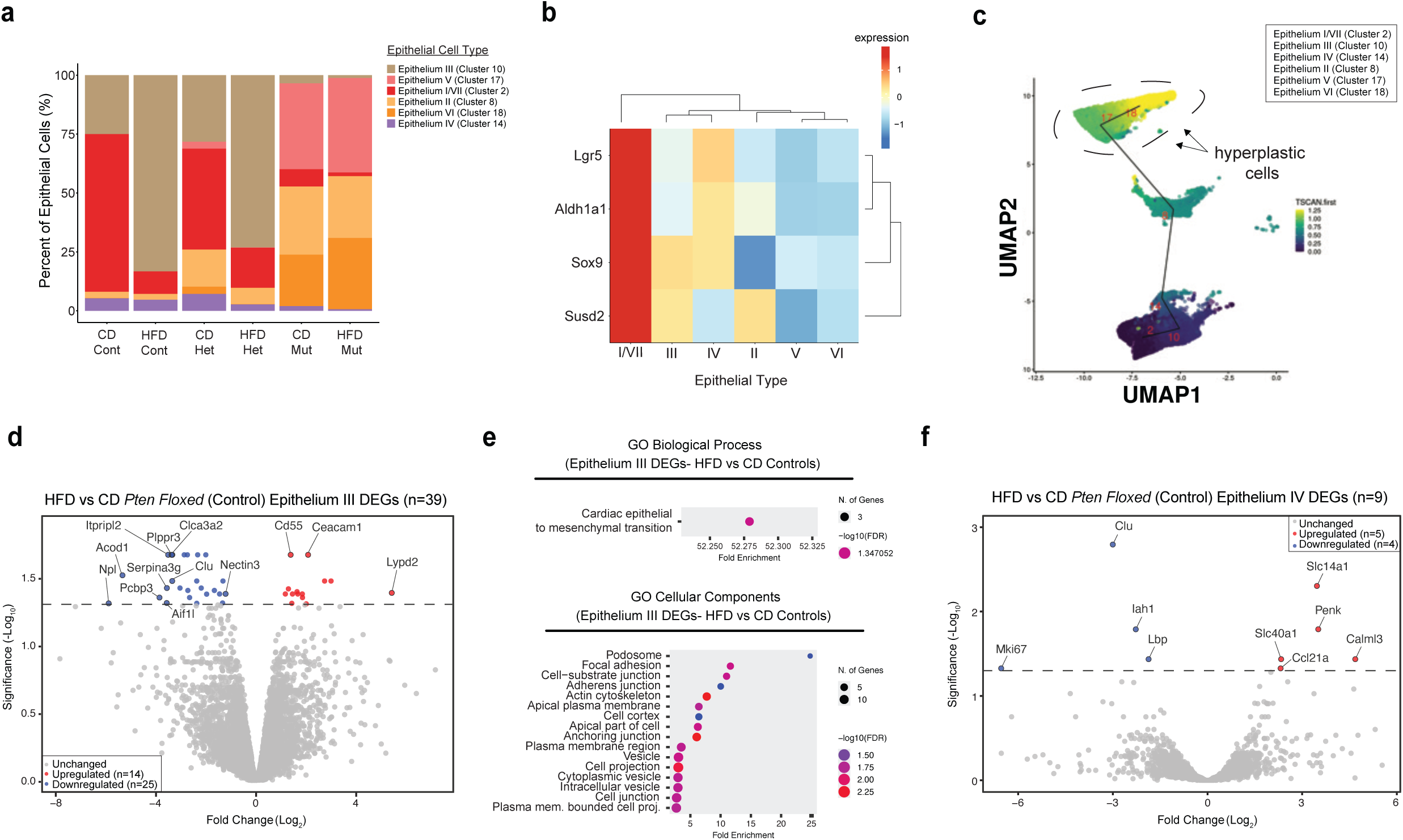
HFD-treated mouse epithelia shift from stem cell–like signatures toward hyperplastic signatures seen in a conditional PTEN-deficient mouse model of EH. (a) Stacked bar graph comparing proportions of the six epithelial cell clusters across CD and HFD conditions with control, *Pten* heterozygous, and *Pten* homozygous mutant conditions. (b) Heatmap of epithelial stem cell marker expression in the six epithelial clusters across all treatment groups. (c) UMAP plot of a pseudotime analysis of the six epithelial clusters from all treatment groups. (d) Volcano plot of HFD versus CD *Pten^floxed^* (control) mouse DEGs (n=39) in Epithelium III. Significantly upregulated (n=14) genes are in red and downregulated (n=25) are in blue. (e) GO Biological Processes and Cellular Components enrichment analyses dotplots from Epithelium III DEGs. (f) Volcano plot of HFD versus CD DEGs (n=9) in Epithelium IV. Significantly upregulated (n=5) genes are in red and downregulated (n=4) are in blue. DEG significance determined at FDR<0.05. Volcano plots were created online with VolcaNoseR^125^. GO enrichment plots were developed online with ShinyGO 0.82^126^.

To better understand the significance of these epithelial subtype shifts, we subsequently performed a cell trajectory analysis on the six epithelial clusters. Epithelium I/VII (Cluster 2) was used as the root for the epithelial clusters due to its expression of proposed endometrial stem cell markers *Lgr5*, *Aldh1a1*, *Sox9*, and *Susd2*^80–82^ (Fig. 6b). Compared to the CD-enriched Epithelium I, the cell trajectory placed the HFD-enriched Epithelium III pseudotemporally closer to the hyperplastic Epithelium V and VI, suggesting that HFD leads epithelium away from stem cell-like signatures towards a cell transitional state closer to hyperplastic epithelium (Fig. 6c). When examined more closely, the Epithelium I/VII stem cell cluster (Fig. 6b) comprised ∼75% of epithelial cells in CD controls (Fig. 6a), a portion that appears unusually high for a stem cell population. Revisiting the UMAP clustering, although Epithelium I constituted the majority of Cluster 2 (Fig. 1d,e; Supplementary Fig. 7a), Epithelium VII was also present in this cluster. This led us to hypothesize that Epithelium VII (within Cluster 2) may represent a more plausible stem cell population within this cluster. We confirmed that Epithelium VII had high expression of stem cell and proliferation (*Mki67*) markers (Supplementary Fig. 7c). Furthermore, Epithelium VII, like Epithelium I, was proportionally reduced in HFD controls (1.82%) compared to CD controls (10.98%) (Supplementary Fig. 7d). When investigating proliferation in the six epithelial cell type clusters, we found that while Epithelium II appeared to have the highest expression levels of *Mki67*, Epithelium I/VII (Cluster 2) had the highest percent of cells with *Mki67* expression, as expected due enriched stem cell marker expression (Supplementary Fig. 7e,f). Across treatment conditions, CD-treated control mice had the highest percent of total epithelia expressing *Mki67* (Supplementary Fig. 7g), which correlates with Epithelium I/VII being the predominant cluster in this treatment group. When stratifying on both diet and epithelial type, the highest percent of *Mki67*^+^ cells in every treatment condition (*Pten*-status and diet) was in Epithelium III (Supplementary Fig. 7h). Cell cycle analysis^83^ was also performed to infer epithelial cell cycle stages by treatment group, epithelial type, or both (Supplementary Fig. 7i-k). PTEN-deficient mice trended toward a higher proportion of epithelia in the G1 phase and a reduced proportion in the S phase.

When examining the epithelial cluster DEGs, only two of the six clusters had HFD-induced DEGs, Epithelium III and IV (Fig. 1f). Epithelium III had the most, with 39 DEGs, of which 14 were upregulated and 25 were downregulated (Fig. 6d). This included an upregulation of *Cd55*, which is a complement-regulatory protein upregulated in EC and cancer stem cells in multiple cancer types^84,85^. CD55 signaling can promote malignant transformation and cancer cell survival, while CD55 inhibition promotes complement-dependent cytotoxicity. Biological Processes enrichment analysis revealed that HFD modulated cardiac epithelial to mesenchymal transition (EMT) (Fig. 6e). Cellular Components GO enrichment indicated alterations to cell junctions, including a downregulation of uterine epithelial cell junction gene, *Nectin3* (Fig. 6l)^86^. EMT, including altered epithelial cell junctions, is associated with tissue fibrosis due to chronic inflammation^87^. The remaining 9 epithelium DEGs were present in Epithelium IV, with 5 upregulated and 4 downregulated genes (Fig. 6f).

### HFD exacerbates EH in a sensitized, *Pten* heterozygous mutant background

In order to further determine if our HFD-induced endometrial changes create conditions favorable to EH development, we utilized our conditional *Pten* heterozygous loss-of-function mice to histologically examine whether these HFD-associated changes persist in the endometrium and thus conceivably worsen EH severity. Non-estrous staged *Pten^fl/+^; Pax8^Cre/+^* heterozygous mice remaining from the scRNA-seq cohort were utilized for histology. Mice were fed either CD or HFD for 26-37 weeks (until 33-43.5 weeks of age) (Supplementary Fig. 1a-d; Supplementary Fig. 8a,b). As seen previously with control animals, HFD *Pten* heterozygous mice displayed an increased final body weight and glucose intolerance, a similar semi-dry uterine weight, and a decreased body-to-uterine weight ratio (Supplementary Fig. 1e-i; Supplementary Fig. 8c,d). We previously demonstrated that estrus-staged HFD-treated mice exhibited reduced circulating progesterone, resulting in an unopposed estrogenic state^28^. Estrus-staged HFD-treated *Pten* heterozygous mice likewise had a reduction in progesterone, trending toward an unopposed estrogen state (Supplementary Fig. 8e-h).

Uterine sections from CD and HFD-treated *Pten* heterozygous mutant mice were stained with H&E or Masson’s Trichrome, and immunostained for cytokeratin 8 (KRT8; epithelium) or PTEN (Fig. 7a-h). Comparative *Pten floxed* control histology is also provided (Supplementary Fig. 9a-f). As expected, based on H&E and KRT8, both CD and HFD-treated *Pten* heterozygous mutants developed EH (Fig. 7a-d) and a selective loss of PTEN in some, but not all, of the glands (Fig. 7e,f). On the other hand, HFD-treated mutant mice displayed increased EH disease severity, as determined by an increased number of glands and increased epithelial height in the abnormal glands, though the percent of normal and abnormal glands, as well as abnormal gland area were similar (Fig. 7i-m). This increased disease severity supports the transcriptomic data from *Pten floxed* (control) samples, suggesting that HFD promotes an epithelial shift toward a state more primed for hyperplastic disease formation. Similar to HFD-treated control animals, HFD-treated *Pten* heterozygous mutants histologically display increased stromal ECM deposition (Masson’s Trichrome) compared to the CD-treated *Pten* heterozygous mutants (Fig. 7g-h,n-q). Increased ECM was present in the endometrium of both estrus-staged and non-estrous staged mice on HFD, thus indicating that this fibrotic stromal phenotype is a persistent versus transient or estrus stage-dependent condition.

**Fig. 7.**
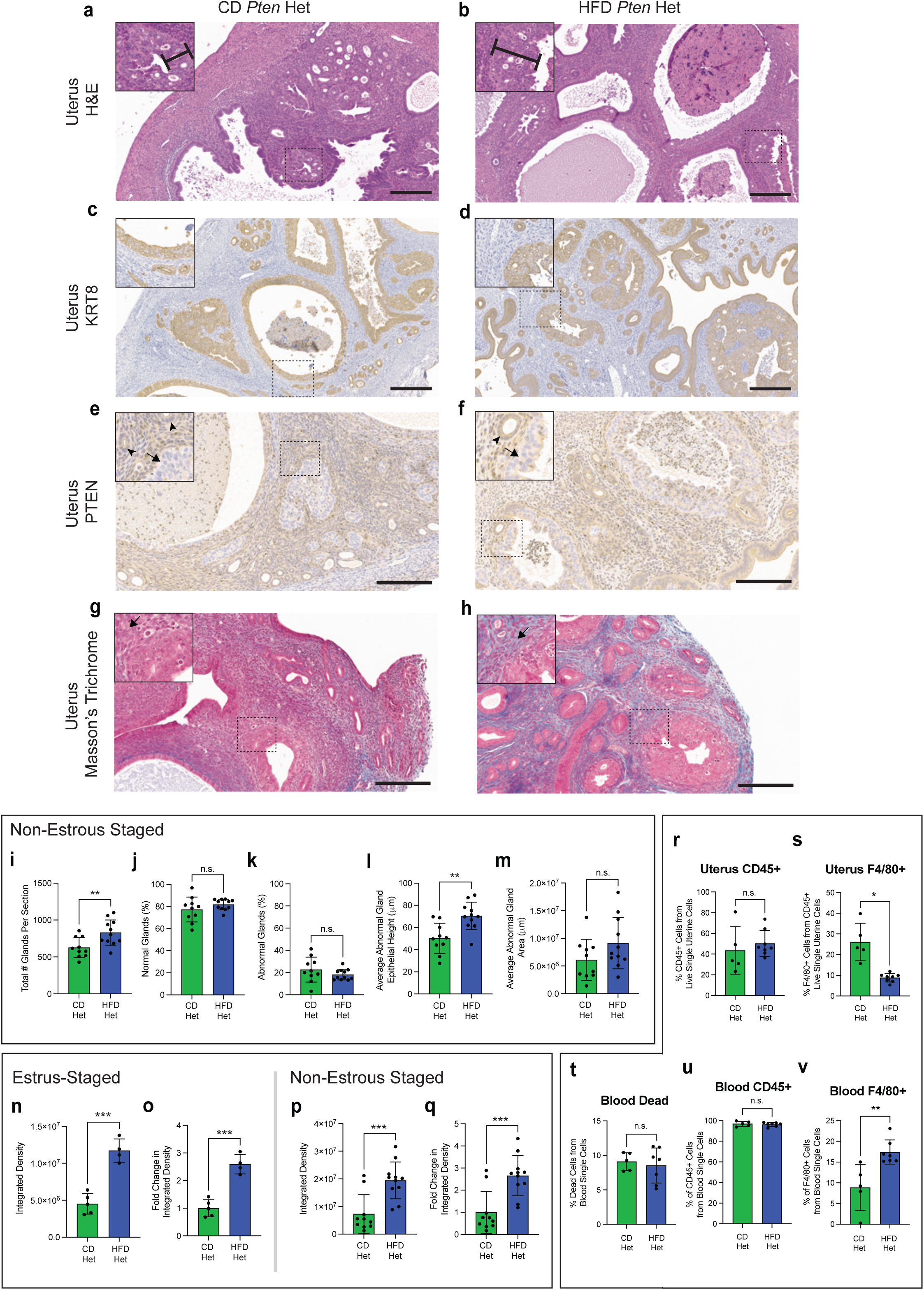
HFD-treated *Pten* heterozygous mutants display worsened EH disease severity, stromal fibrosis and a reduction in macrophages in the uterus. (a,b) Representative H&E stain of endometrial tissue from *Pten* heterozygous mutants for each diet. Main image= 10x magnification; inset=20x magnification; scalebar=300 μm. Caliper indicates the height of an abnormal endometrial gland. (c,d) Representative KRT8 IHC stain of endometrial tissue from *Pten* heterozygous mutants for each diet (CD n=5; HFD n=4). Main image= 10x magnification; inset=20x magnification; scalebar=300 μm. (e,f) Representative PTEN IHC stain of endometrial tissue from *Pten* heterozygous mutants for each diet (CD n=9; HFD n=11). Main image= 20x magnification; inset=40x magnification; scalebar=200 μm. Arrow indicates a gland with PTEN loss, while an arrowhead indicates a PTEN intact gland. (g,h) Representative Masson’s Trichrome stain of endometrial stromal ECM from *Pten* heterozygous mutants for each diet. Main image= 20x magnification; inset=40x magnification; scalebar=200 μm. Arrow indicates endometrial stroma. (i-m) Graphs quantifying EH disease severity in either CD- (n=10) or HFD- (n=11) treated non-estrous staged *Pten* heterozygous mutant mice from H&E-stained endometrial slides. Measured disease severity parameters were total number of glands per representative histological section, percent normal glands (%), percent abnormal glands (%), average abnormal gland epithelial height (μm), and average abnormal gland area (μm). Datapoints represent a value for each mouse (bar=mean ± s.d.). Two-tailed unpaired *t*-test statistic with Welch’s correction as needed. (n-q) Graphs quantifying ECM raw integrated density or fold change in integrated density in *Pten* heterozygous mutant endometrial stroma using Image J with the Color Deconvolution plugin. Samples were from either estrus-staged or non-estrous staged mice in both CD- (estrus n=5; non-estrus n=10) or HFD- (estrus n=4; non-estrus n=11) treated animals. Measurement (datapoint) represents each mouse’s mean of 4 samplings (bar=mean ± s.d.). Two-tailed unpaired *t*-test statistic. (r) Percent (%) of CD45^+^ cells from all live single cells in the uterus of CD (n=5) and HFD (n=8) *Pten* heterozygous mice. Measurement (datapoint) represents each mouse (bar=mean ± s.d.). Two-tailed unpaired *t*-test statistic. (s) Percent (%) of F4/80^+^ cells from all CD45^+^ live single cells in the uterus. Measurement (datapoint) represents each mouse (bar=mean ± s.d.). Two-tailed unpaired *t*-test statistic with Welch’s correction. (t,u) Percent (%) of dead cells or CD45^+^ immune cells from all single cells in blood of CD (n=5) and HFD (n=7) *Pten* heterozygous mice. Measurement (datapoint) represents each mouse (bar=mean ± s.d.). Two-tailed unpaired *t*-test statistic. (v) Percent (%) of F4/80^+^ cells from CD45^+^ single cells in the blood. Blood F4/80^+^ cells are likely monocytes, as macrophages are not commonly seen in circulation. Measurement (datapoint) represents each mouse (bar=mean ± s.d.). Two-tailed unpaired *t*-test statistic. *\*p<0.05; **p<0.01; ***p<0.001; ****p<0.0001*

### F4/80^+^ monocytes are increased in HFD circulation, but F4/80^+^ macrophages are reduced in HFD endometrium

In addition to ECM-related genes, fibroblast DEGs indicated alterations to the immune system including reduced monocyte recruitment and maturation. Circulating monocytes migrate to target tissues where they mature into macrophages^74^. Accordingly, a reduced proportion of macrophages was observed in the HFD-treated endometrium, which may contribute to the unresolved ECM accumulation and persistence of abnormal epithelial cells. To verify this reduction and compare levels to those in circulation, we placed an additional cohort of conditional *Pten* heterozygous loss-of-function mice on either CD or HFD for 21–30 weeks (27.5-38 weeks of age) (Supplementary Fig. 8i-n). We then collected estrus-staged uteri and blood for macrophage flow cytometry analysis (Supplementary Fig. 8o,p). After live single cell enrichment, there was no difference between CD and HFD in the percent of CD45^+^ immune cells (out of all live uterine cells), but there was a significant decrease in the percent of F4/80^+^ macrophages (out of all CD45^+^ cells) (Fig. 7r,s). As expected, after red blood cell removal nearly all the cells from the blood were CD45^+^ immune cells (all were over 92%) and there was no difference between diets in the percent of dead cells (Fig. 7t,u). In contrast to the uterus, the percent of blood F4/80^+^ cells was increased with HFD (Fig. 7v). These blood F4/80^+^ cells are likely monocytes, as monocytes can express F4/80, albeit at a lower level than macrophages, and macrophages are rarely in circulation^74,88^. Subsequent analysis was performed to further characterize these F4/80^+^ cells (Supplementary Fig. 8q). Pro-inflammatory macrophages, which we characterized as CD80^+^, are thought to be dominant in the ovary, while anti-inflammatory macrophages, which we labeled as CD206^+^, are dominant in the uterus^38^. While HFD-treated blood had an increase in pro-inflammatory CD80^+^ F4/80^+^ cells, there was no difference in the uterus (Supplementary Fig. 8r,u). There was no significant difference in anti-inflammatory CD206^+^ F4/80^+^ cells in the HFD-treated uterus or blood (Supplementary Fig. 8s,v). There was also no significant difference when comparing the ratio of CD80^+^ F4/80^+^ to CD206^+^ F4/80^+^ cells in the HFD-treated blood or uterus (Supplementary Fig. 8t,w). Overall, while HFD is associated with a higher percent of circulating F4/80^+^ monocytes, there was a decrease in F4/80^+^ macrophages in the uterus, which was consistent with our scRNA-sequencing data in *Pten floxed* controls. Thus, the HFD-associated reduction in uterine macrophages may impair endometrial stromal repair and permit the persistence of abnormal epithelial cells.

## Discussion

Endometrial homeostasis is tightly regulated by cycle-dependent fluctuations in estrogen and progesterone^1^. Obesity disrupts this hormonal balance by decreasing progesterone production, thereby promoting a comparatively unopposed estrogen state^26^. Since progesterone counteracts the pro-proliferative effects of estrogen on the epithelium, this imbalance is associated with an increased risk of EH development and greater disease severity^18–20,89^. While prior studies utilizing the HFD mouse model show systemic signs of obesity, they do not exhibit histological evidence of EH^28,29^. Here, we employed a known genetically induced mouse model of EH in combination with scRNA-seq to better elucidate the cell-type-specific effects of obesity on the initiation and progression of endometrial disease. We reveal that HFD disrupts endometrial homeostasis by transcriptomically reprogramming fibroblasts toward inflammatory and ECM-enriched states. These transcriptomic changes are consistent with our histological findings of stromal ECM accumulation, indicative of fibrosis or unresolved tissue damaged driven by persistent inflammation. Using a conditional *Esr1* mutant mouse model, we verified that stromal estrogen signaling mediates HFD-induced ECM accumulation. Consistent with estrogen-driven stromal pathology, HFD reduced endometrial PGR-network-associated Aldh1a2^+^ Fibroblasts, thus proportionally expanding inflammatory ER-network-associated Gsn^+^ Fibroblasts. Downregulation of monocyte and macrophage recruitment and activation genes corresponded with decreased uterine macrophages, likely contributing to the persistent ECM accumulation and fibrosis. HFD additionally shifted the predicted epithelial cell lineage towards EH cells, suggesting that HFD predisposes the epithelium to hyperplastic disease. Moreover, when combining HFD and heterozygous PTEN loss, HFD exacerbates EH disease severity. Like the HFD-treated control animals, stromal fibrosis and uterine macrophage depletion persisted in HFD mice genetically predisposed to EH, suggesting that HFD-induced dysregulation in the mutation-free fibroblasts are implicated in EH disease development and severity.

HFD-induced obese mice have unopposed estrogen levels, due to diminished post-ovulatory progesterone^28^. Endometrial estrogen signaling indirectly induces epithelial proliferation though the endometrial fibroblasts^6–9^, underscoring the importance of stromal homeostasis on endometrial function. In our HFD-treated mice, fibroblasts had the highest number of DEGs. Among the fibroblast subtypes, we observed a proportional expansion of ER-network-associated Gsn^+^ Fibroblasts due to a reduction in the PR-network-associated Aldh1a2^+^ Fibroblasts. PGR protein expression is reduced in the endometrium of women with obesity^90^. Gsn^+^ Fibroblast marker genes, *Clec3b*, *Col6a3*, *Col14a1*, *Dpt*, and *Efemp1*, are observed in fibroblasts found throughout the murine endometrium that are enriched for ECM production and collagen deposition^91^. We found the Gsn^+^ Fibroblasts to be distributed throughout the endometrium in both CD- and HFD-treated conditions, though appeared more prominent in the HFD-treated samples. This correlates with scRNA-seq data indicating a proportional increase in Gsn^+^ Fibroblasts, as well as an upregulation of *Gsn* and ECM genes with HFD. Almost half (n=23/50) of Gsn^+^ Fibroblast I marker genes are shared with a population of human endometrial “C7 fibroblasts”, including *Cxcl12*, *Dcn*, *Dpt*, *Gsn*, *Lum*, *Mgp*, *Serping1*, and *Timp2*^42^. *C7* was similarly upregulated in our HFD-treated Gsn^+^ Fibroblast population. C7 fibroblasts are known to reside in the basal layer of the human endometrium throughout the menstrual cycle^92^ and communicate broadly with a wide range of endometrial stroma, immune, epithelium, and endothelium subtypes^44,93^. Notably, one cell-cell interaction involves CXCL12 produced by C7 fibroblasts signaling to CXCR4 on SOX9/CHD2^+^ basalis endometrial epithelial cells, which are predicted to function as epithelial stem cells^94^. CXCL12 can recruit bone marrow-derived stem cells to help repair fibrotic endometrial tissue in Asherman’s syndrome, which is particularly important given the potential depletion of endogenous stem cells caused by damage to the basal layer of the endometrium^95^. In contrast, the Aldh1a2*^+^* Fibroblast subtype is predicted to be a heterogeneous population of sub-epithelial and “inner” fibroblasts in the murine endometrium due to enrichment of *Aldh1a2*, *Angptl7*, *Cdh11*, *Cxcl14*, *Fn1*, *Islr*, *Ngfr*, *Slc26a7*, and *Smoc2*^91^. Functionally, sub-epithelial or “inner” fibroblasts are important for response to wounding and regulation of the immune response.

In regard to the estrous cycle, Aldh1a2^+^ Fibroblast marker genes, *A2m*, *Aldh1a2*, *Ccl11*, *Col6a4*, *Dio2*, *Hsd11b2*, *Htra1*, and *Smoc2*, are enriched in mouse endometrial maturational fibroblasts associated with the second half of the estrous cycle or the secretory phase in humans^36^. The Aldh1a2^+^ Fibroblast markers *Aldh1a2*, *Cdh11*, *Dio2*, *Islr*, *Mdk*, and *Nid1* are also potential markers for human decidualized fibroblasts^44^. In humans, the progesterone-dominated secretory phase induces fibroblasts to differentiate into decidual cells that drive ECM remodeling, modulate the immune response, and enhance resistance to oxidative stress—collectively preparing the endometrium for successful embryo implantation^96^. Aldh1a2^+^ Fibroblast II displayed HFD-induced enrichment for mitochondrial depolarization, which can be caused by obesity-induced oxidative stress within the cell^55^. Correspondingly, *Sod3*, which is thought to protect against HFD-induced oxidative stress^57^, was upregulated in all three fibroblast subtypes. SOD3 overexpression significantly decreases monocyte recruitment to sites of tissue damage^97^.

Inflammatory pathways are most prominent in endometrial fibroblasts in the estrogen-dominant phases of the cycle^38^. Conversely, fibroblasts from progesterone-dominant phases of the estrous cycle promote anti-inflammatory signaling. Fibroblasts are important regulators of the immune response, from sustaining chronic inflammation to promoting immunosuppression in cancer^73^. In the murine uterus, spatial transcriptomic analyses showed that autocrine endometrial fibroblast signaling more likely controlled tissue inflammation status rather than immune cells^38^. In our study, HFD upregulated immune system components in the dominant ER-network-associated Gsn^+^ Fibroblast cluster, with the majority involving the classical complement pathway. *C3* is one of the upregulated complement factors observed in our dataset. In the murine endometrium, estrogen stimulates *C3* expression^98^, whereas progesterone inhibits its expression^99^. Excessive complement activation can lead to uncontrolled inflammation and tissue damage^45,46^.

Therefore, estrogen dominance likely contributes to a chronic proinflammatory state through complement production, leading to unrepaired tissue damage. In the context of tissue damage, ECM-related genes were also greatly enriched in this subtype and congruently, HFD-stained endometrium displayed increased stromal ECM deposition, indicative of fibrosis or unresolved scarring. Sustained inflammation can drive tissue fibrosis and the ensuing excessive ECM remodeling can impede immune cell infiltration, thereby limiting the clearance of abnormal cells^38^. By using an additional ERα (*Esr1*) cKO mouse model, we further confirmed that stromal estrogen signaling was implicated in the HFD-induced endometrial ECM accumulation. This suggests that prolonged estrogen dominance, due to diminished post-ovulatory progesterone, induces a chronic proinflammatory state, which disrupts endometrial fibroblast homeostasis and generates tissue fibrosis with HFD.

Macrophages are fundamental for clearing abnormal cells and debris, regulating inflammation, and promoting repair of the endometrium^5,75^. A reduction in endometrial macrophages has also been linked to the development of Asherman’s syndrome, a trauma-induced fibrotic disease of the endometrium^75^. The majority of uterine macrophages are cyclically replenished from circulating bone marrow-derived monocytes^100,101^. Although circulating monocytes were elevated with HFD, uterine macrophages were decreased. This reduction may result from both physical barriers imposed by dense ECM^102^, as well as altered fibroblast and monocyte gene expression that collectively inhibit immune cell maturation and recruitment. NK cells are commonly studied in the context of pregnancy, where these cells play a crucial role in establishing immune tolerance in the uterus and function as immunosuppressive cells^76^. HFD likely kept NK cells in a resting state, as there was a downregulation of the active-state associated *Gzmc*. This inactive state was likely influenced by fibroblasts, as NK cells are activated by progesterone-induced IL-15 production in fibroblasts^53,78^, and *Il15* expression was downregulated in our HFD-treated Gsn^+^ Fibroblasts. Thus, Gsn^+^ Fibroblasts may also play an important role in NK cell development and activation in the progesterone dominant phase of the cycle and pregnancy. Collectively, HFD-induced dysregulation of monocytes, macrophages, and NK cells may allow persistent inflammation and excessive ECM accumulation in the endometrial stroma.

Estrogen-dominance and stromal fibroblast and immune cell transcriptomes suggest an environment conducive to epithelial proliferation and impaired clearance. Even though histologically evident EH formation was absent with HFD, the HFD-dependent shift from a stem cell-like signature towards a transitional state closer to hyperplastic epithelium was evident. Epithelium III was the predominant epithelial type in HFD-treated control endometrium. When examining Epithelium III DEGs, *Cd55* was upregulated. CD55 is upregulated in EC and functions to promote malignant transformation and cancer cell survival^84,85^. Therefore, upregulation of *Cd55* supports the shift of HFD epithelium to a transitional state closer to hyperplastic epithelium.

Not only does obesity increase a woman’s risk of developing EH, obese women with EH are also more likely to develop more serious or progressive forms of the disease, such as complex EH and EH with atypia, which have a higher potential to further progress into EC^103^. This raised a question: if EH is already present, do HFD-induced fibrotic stroma and decreased macrophages persist in the endometrium, and therefore have the potential to contribute to increased disease severity associated with obesity? With the addition of conditional *Pten* mutation to genetically induce EH, HFD led to worsened EH severity compared to CD, and the stromal ECM accumulation and decreased uterine macrophages were again present. Therefore, HFD-associated fibrosis and dampened macrophage recruitment may drive the HFD endometrium both towards development and increased progression of EH.

In addition to obesity, perimenopause and advancing age increases EH risk^104,105^. Risk increases at this age as perimenopause is characterized by irregular menstrual cycles, which can include episodes of elevated estrogen and anovulation, leading to low progesterone^106^. Aging is also associated with an increase in endometrial fibroblast inflammation and uterine fibrosis^38^. Therefore, therapies that counteract the effects of unopposed estrogen on endometrial fibroblasts may be critical for preventing inflammation-driven endometrial fibrosis and reducing the risk of creating an environment that promotes the development and progression of EH, not only in women with obesity but also more broadly across the aging female population.

In summary, chronic HFD-induced estrogen dominance disrupts endometrial fibroblast homeostasis to induce persistent inflammation and ECM accumulation, indicative of fibrosis and sustained tissue damage. We hypothesize the resulting stromal dysfunction impairs immune cell infiltration and function and fosters an environment permissive to unchecked epithelial proliferation, impairing endometrial function and collectively promoting the development and progression of EH.

## Methods

### Mice

*Pten ^flox^* (RRID:IMSR_JAX:006440)^107^, *Pax8^cre^* (RRID:IMSR_JAX:028196)^108^, and *Esr1^flox^* (RRID: IMSR_JAX:032173)^68^ alleles were purchased from The Jackson Laboratory and the *Amhr2^Cre^* (RRID:MGI:3042214)^67^ allele were obtained from Drs. Tae Hoon Kim and Jae-Wook Jeong at the University of Missouri. Mice were then further bred and crossed in-house at Michigan State University’s (MSU) Grand Rapids Research Center (GRRC) vivarium. Age- and litter-matched *Pten ^floxed^* (control), *Pten^fl/+^;Pax8^Cre/+^*, and *Pten^fl/fl^;Pax8^Cre/+^* or *Esr1^floxed^;Amhr2^+/+^* and *Esr1^fl/fl^;Amhr2^Cre/+^*mice were placed on CD or HFD for a minimum of 21.5 weeks female mice on a C57BL/6J background were assigned to either a 60% high-fat diet (HFD) (Research Diets, D12459) or a matched low-fat control diet (CD) treatment condition (Research Diets, D12450J). Mice were placed on diet starting at 5-12-week-old in all cohorts. Similar to our previous study, we performed weekly cage changes and recorded mouse weight along with the amount of food eaten per cage^28^. Mice were fed *ad libitum* for 30-35 weeks in the *Pten ^floxed^* (control) scRNA-seq cohort (35-43 weeks of age), 26-37.5 weeks in the *Pten^floxed^;Pax8^Cre/+^*(heterozygous and homozygous mutant) histology and scRNA-seq cohort (27.5-43.5 weeks of age), 21–30 weeks in the macrophage flow cytometry cohort (27.5-38 weeks of age), and 21.5-31.5 weeks in the *Esr1^floxed^* cohort (28-38.5 weeks of age). Mice were tested for glucose intolerance at 17.5-25 weeks on diet, with a minimum of two weeks for recovery before tissue harvest. Mice were then euthanized and tissue collected during the estrus-stage of the estrous cycle (unless otherwise noted), as determined by vaginal lavage and cytology as previously described^28^. Blood was collected at tissue harvest via a cheek bleed for later serum hormone levels or immune cell flow cytometry analysis. Mice were then euthanized by CO_2_ inhalation. Uterus, ovaries, adipose tissue, liver, kidney, spleen, and thyroid were collected and placed in 10% neutral-buffered formalin (NBF) (Epredia, Ref#9400). For mice used in scRNA-seq and macrophage flow cytometry, a small portion of a uterine horn was saved for fixation, as most of the uterus was allocated for scRNA-seq and flow cytometry respectively. Mice were housed on a 24-hour light cycle where light-phase was 7 AM (Zeitgeber time; ZT0) to 7 PM (ZT12) and dark-phase was 7 PM (ZT12) to 7 AM (ZT0). Mice were housed in the vivarium at MSU’s GRRC in agreement to approved protocols from MSU’s Institutional Animal Care and Use Committee (IACUC). MSU has an approved Animal Welfare Assurance from the National Institutes of Health (NIH) Office of Laboratory Animal Welfare (OLAW) and is accredited by the Association for Assessment and Accreditation of Laboratory Animal Care (AAALAC). MSU is registered with the US Department of Agriculture (USDA).

### Glucose Tolerance Test

Glucose tolerance test was performed as previously described^28^. In summary, after 17.5-25 weeks all mice were tested for glucose intolerance. At 8 AM (ZT1), mice weighed and then fasted for six hours with free access to water. After six hours of fasting (ZT7), the tip of the mouse’s tail was cut, and a drop of blood was collected to determine fasting glucose levels with a glucometer. Based on body weight, the mouse was then injected with autoclaved USP D(+)-Glucose dissolved in 0.9% saline. A single drop of blood was then collected additionally at 30-, 90-, 150-, and 210-minutes after dextrose injection.

### Single-Cell RNA Isolation for 10x Genomics’ Chromium Single Cell Gene Expression Flex Protocol

After 30-35 weeks on CD and HFD, Pten*^floxed^* (control) estrus-staged mice were euthanized and uteri collected. CD and HFD *Pten^fl/+^;Pax8^Cre/+^* (after 26-37 weeks on diet) and *Pten^fl/fl^;Pax8^Cre/+^* mice (after 23-32 weeks on diet) with EH were additionally collected for additional comparison of control mice to a diseased state. After dissection, uteri were minced with scissors and then digested enzymatically at 37°C using a Multi Tissue Dissociation Kit 2 enzyme cocktail (Miltenyi Biotec, cat# 130-110-203) and mechanically using the gentleMACS™ Octo Dissociator. Digested tissues were then poured through a 40 μm sterile filter (Fisher Scientific, cat#22-363-547) and then red blood cells lysed (Miltenyi Biotec, cat#130-094-183). Cells were then sorted to isolate live cells using the magnetic assisted cell sorting (MACS) Dead Cell Removal Kit (Miltenyi Biotec, cat#130-090-101). A minimum of 300,000 cells per sample was then aliquoted and filtered with a Flowmi™ 40 μm tip strainer (SP Bel-Art, cat#H13680-0040). Multiple isolation days were needed to collect uteri in estrus, therefore cells were processed for 10x Genomics’ Chromium Single Cell Gene Expression Flex (v1) protocol, which is a probe-based gene expression design and allows for cells to be fixed and stored frozen until quality control (QC) and subsequent probe hybridization. Storing the cells was preferred to avoid batch effect of different library preparation and sequencing runs. Cells were fixed using Chromium Next GEM Single Cell Fixed RNA Sample Preparation Kit (10x Genomics, cat#PN-1000414) and cold Formaldehyde with Fix & Perm Buffer (10x Genomics, cat#PN-2000517) and left overnight at 4°C. The next day, cells were resuspended in cold Quench Buffer (10x Genomics, cat#PN-2000516), counted, and Enhancer (10x Genomics, cat#PN-2000482) with glycerol added. Cells were then stored at -80°C until all samples were collected.

### Construction and Sequencing of 10X FLEX Libraries

Libraries were generated and sequenced by the Van Andel Institute Genomics Core (Grand Rapids, MI, USA). Before probe hybridization, the quality of the samples was assessed using the Luna-FX7 (Logos Biosystems, Dongan-gu Anyang-si, Gyeonggi-do, South Korea), with additional image analysis to quantify cell aggregates, debris, and cell number using image software Cell Profiler (Broad Institute, Cambridge, MA, USA). Samples that met the quality and quantity criteria provided by 10X Genomics, moved into the 10X (v1) Gene Expression kit Next GEM with in-line multiplexing (10X Genomics, Pleasanton, CA, USA) workflow using the Chromium Fixed RNA kit, Mouse transcriptome, 4rxns x 4BC (PN-1000496) according to the manufacturer’s instruction to target an output of 10,000 cells per sample using a 10X Genomics ChromiumX Controller. Briefly, samples were hybridized to one of 4 uniquely barcoded probe sets, allowing for multiplexing up to 4 samples. Each gene target contains 2 probes designed to bind both sides of the gene target, which are ligated during GEM incubation to ensure target specificity. After hybridization, the samples were again imaged on the Luna-FX7 (Logos Biosystems, Dongan-gu Anyang-si, Gyeonggi-do, South Korea) and pooled in equal ratios with each other. The pooled samples were washed several times and filtered twice with a 30 µm Pre-Separation filter (Miltenyi Biotec, Bergishch Gladbach, German). Single cells were captured in gel beads in emulsion (GEMs), which, after GEM incubation, were lysed, releasing uniquely barcoded probes containing barcodes for both the individual cell, the sample identifier, and unique molecular identifiers. The ligated product is then pre-amplified before undergoing another round of PCR to add P5, P7, i5, and i7 sample indexes and the Illumina TruSeq Read 1 sequence (Read1T) and Small Read 2 (Read 2S) sequences. Quality and quantity of the finished gene expression libraries were assessed using a combination of Agilent DNA High Sensitivity chip (Agilent Technologies, Inc.) and QuantiFluor® dsDNA System (Promega Corp., Madison, WI, USA). 28x10x10x90 bp, paired end sequencing was performed on an Illumina NovaSeq 6000 sequencer (Illumina Inc., San Diego, CA, USA) using S1, 100 cycle sequencing kit (v1.5) to a minimum raw read depth of 10K reads per cell. Base calling was done by Illumina RTA3 and output was demultiplexed and converted to FastQ format with Illumina Bcl2fastq v2.20.0.

### Single-Cell RNA Sequencing Analysis

scRNA-sequencing data analysis workflow was performed by the Bioinformatics & Biostatistics Core at the Van Andel Institute (Grand Rapids, MI, USA). Fastq files were aligned using the ‘multi’ workflow from Cellranger v7.1.0 with the ‘refdata-gex-mm10-2020-A’ reference and the ‘Chromium_Mouse_Transcriptome_Probe_Set_v1.0.1_mm10-2020-A.csv’ probeset file provided by 10x. The filtered per-sample gene counts were processed using an R/Bioconductor workflow that encompasses normalization, batch correction, differential expression between conditions within clusters, and marker gene detection between clusters^109^. Using the scuttle v1.8.4 package^110^, total UMI count, number of genes detected, and percent of counts from mitochondrially-encoded genes were calculated, and used to filter low-quality cells based on cutoffs calculated on a per-sample basis. Filter thresholds for total UMI count and number of genes detected were calculated using the perCellQCFilters function with the default cutoff of 3 median absolute deviations, on the log scale; the threshold for percent mitochondria was calculated by calling the isOutlier function directly, setting ‘nmads = 5’ and ‘type = higher’. For doublet detection, each sample was processed separately. Normalized counts were calculated using the ‘deconvolution’ method, which involves pooling cells to overcome the large number of genes with zero counts in individual cells, enabling the use of bulk RNA-seq normalization approaches^111–113^. Specifically, the quickCluster, computeSumFactors and logNormCounts functions from scran v1.26.2 and scuttle v1.8.4 were used with default settings^114^. After normalization, the top 3000 most variable genes were identified using the getTopHVG function and used for Principal Component Analysis (PCA) with the fixedPCA function. Clusters were identified using the clusterCells function with default settings. Doublet scores were calculated by running the computeDoubletDensity function with ‘d = 50’ and the scDblFinder function with default parameters, both of which two separate methods from scDblFinder v1.12.0 were used to calculate doublet scores, both of which simulate doublets by combining pairs of randomly chosen cells but have distinct scoring schemes^115^. These doublet scores were visualized with cluster annotations to assess the proportion of likely doublets in each cluster. To reduce technical variation between samples that can diminish the ability to cluster together similar cell types, the quickCorrect function from batchelor v1.14.1 was run with the parameter, ‘PARAM = FastMnnParam(auto.merge = TRUE)’, batch-correcting the PCA coordinates using the mutual nearest neighbors (MNN) approach^116^. The corrected coordinates were used for calculating UMAP embeddings using the runUMAP function. In addition, the MNN-corrected coordinates were used to cluster the cells by running the Louvain community detection algorithm on a shared nearest neighbor (SNN) graph; specifically, the clusterCells function, from bluster v1.8.0, was run with the parameter, “SNNGraphParam(k = snn_k, cluster.fun = ‘louvain’)”, where ‘k’ sets the number of nearest neighbors and ‘snn_k ‘ represents different values used to obtain clusters at different resolutions. Sub-clustering of cluster 5 was performed by subsetting the dataset to just the cluster 5 cells, and repeating the quickCorrect and clusterCells functions as described above.

Automated cell type prediction was conducted using SingleR v2.0.0, which compares expression profiles between individual cells in a query dataset and a labeled reference dataset by utilizing Spearman correlations^117^. In the initial run of SingleR, the ‘label.main’ labels in the built-in mouse dataset from the SingleR companion package, celldex v1.8.0, consisting of 358 bulk RNA-seq samples of sorted cells, was used^118^. In a separate run of SingleR, the ‘Idents.pbmc.’ labels from the adult uterus dataset in the mouse cell atlas (MCA) 2.0 were used^35^. For the celldex reference, the pre-calculated log normalized counts were used, while the logNormCounts function was used to calculate log normalized counts for the MCA reference. For the MCA reference, the ‘de.method’ was set to ‘wilcox’, as suggested by the SingleR authors, because the default ‘de.method’ can perform poorly for single-cell reference datasets with many zero gene counts. Downstream processing utilized the ‘pruned.labels’ from SingleR, where low-confidence label assignments were converted to NA. Labels with less than 100 cells were renamed to “Others”, and NA was converted to “No_annot”.

Differential expression analysis between diet-genotype combinations within a given cluster or cell type was conducted using a ‘pseudobulk’ approach with the edgeR quasi-likelihood workflow as implemented in scran v1.26.2^114,119^. The raw counts for the cells from the same sample and cell type or cluster were summed using the aggregateAcrossCells function from scuttle v1.8.4, excluding pseudobulks consisting of <10 cells. For each cluster, diet-genotype groups with <2 pseudobulks were removed, and the pseudoBulkDGE function from scran was used to fit the design, “∼0 + Group”, where Group represented the diet-genotype combinations, and test each contrast. A significance cutoff of 0.05 adjusted P-value was used. Similarly, differential abundance of cells for each cluster was tested using the edgeR quasi-likelihood workflow as described in the “Orchestrating Single-Cell Analysis with Bioconductor” book^109^. The model design was again “∼0 + Group”. The estimateDisp function was run with the parameter, “trend = none”, and glmQLFit was run with “robust = TRUE, abundance.trend = FALSE”. Specific contrasts were tested using the glmQLFTest function.

Per-cell activity scores for transcription factors^120^ were calculated using the run_mlm function with “.mor = weight” and the run_ulm function, respectively, from decoupleR v2.12.0^121^. Activity scores were compared between clusters using Wilcoxon tests, blocking by sample, as implemented in the findMarkers function in scran v1.34 and using the parameter, “lfc = 0.5”, to set the minimum difference between groups. Significance set at: y ≤ 0.0001 (****), y ≤ 0.001 (***), y ≤ 0.01 (**), y ≤ 0.05 (*).

Pseudotime analysis was conducted for the epithelial clusters by constructing a minimum spanning tree (MST) on the cluster centroids using the corrected PCA coordinates and the quickPseudotime function from TSCAN^122^. TSCAN v1.36 was used with the parameter, “outgroup=TRUE” so that distant clusters, if present, can be placed in separate trajectories.

Cell cycle stages were inferred using the tricycle method^83^ as implemented in tricycle v1.6.0. Log-normalized counts were used to project the cells to a reference mouse cell cycle embedding, provided by the tricycle package, using the project_cycle_space function with ‘species = mouse’. Then, “cell cycle position” was estimated using the estimate_cycle_position function, and used to assign each cell as S (0.5 × 𝜋 ≤ 𝑝𝑜𝑠𝑖𝑡𝑖𝑜𝑛 < 𝜋), G2M (𝜋 ≤ 𝑝𝑜𝑠𝑖𝑡𝑖𝑜𝑛 < 1.75 × 𝜋) or G1 (𝑝𝑜𝑠𝑖𝑡𝑖𝑜𝑛 ≥ 1.75 × 𝜋 | 𝑝𝑜𝑠𝑖𝑡𝑖𝑜𝑛 < 0.5 × 𝜋), using the position bins suggested in the package vignette.

The SingleCellExperiment object was split by genotype, removing doublet clusters, then converted into separate Seurat objects using the as.Seurat function. Markers for each cluster were identified using the FindAllMarkers function from Seurat, with the parameters, ‘assay="originalexp", only.pos = TRUE, logfc.threshold=1’. Marker genes and cluster annotations were compiled into CyteType format using the PrepareCyteTypeR function, and submitted to their server using the CyteTypeR function, with the parameter ‘study_context = "Mouse uterine cells"’. Top 50 marker genes for fibroblast clusters were obtained from the online CyteType report^123^. Exploration of results was aided by a ShinyCell instance generated using ShinyCell v2.1.0^124^. Volcano plots were created online with VolcaNoseR^125^. GO enrichment plots were developed online with ShinyGO 0.82^126^.

### Serum Hormone Levels

Mice were in the estrus stage when blood was collected by cheek bleed at the time of tissue harvest. Blood was further processed and analyzed as previously described^28^. In short, blood was mixed with inhibitors, left to coagulate, and then centrifuged and subsequent serum supernatant collected and stored at -80°C. Samples were then shipped to the Wisconsin National Primate Research Center’s Assay Services Unit at the University of Wisconsin-Madison where serum estrone, 17β-estradiol, and progesterone levels were measured using liquid chromatography with tandem mass spectrometry.

### Histology and Immunohistochemistry

NBF-fixed uterine and adipose tissues were paraffin-embedded and sectioned by the Pathology and Biorepository Core at the Van Andel Institute (Grand Rapids, MI, USA). Slides were then stained with routine Hematoxylin and Eosin (H&E), Masson’s Trichrome Aniline Blue Stain Kit (Newcomer Supply, Ref#9179), or Picrosirius Red Stain Kit (Polysciences, Ref#24901) according to the manufacturers’ instructions. Unstained uterine and adipose tissue sections were then stained for various markers using indirect immunohistochemistry (IHC). For IHC, deparaffinized sections underwent heat-based antigen retrieval with either 10mM sodium citrate (pH 6.0) or Tris-EDTA (pH 9.0). Sections were then incubated at the following dilutions: 1:600 Gelsolin (D9W8Y) (Cell Signaling, cat#12953, RRID: RRID:AB_2632961), 1:100 KRT8 (Developmental Sciences Hybridoma Bank, cat#TROMA-I, RRID: AB_531826), 1:200 PTEN (138G6) (Cell Signaling, cat#9559, RRID: AB_390810), and 1:200 ER-a (D6R2W) (Cell Signaling, cat#13258, RRID: AB_2632959). After overnight incubation, biotin-conjugated secondary antibodies were added at the following dilutions: 1:250 donkey anti-rabbit IgG (Jackson Immuno-Research Lab, cat#711-065-152, RRID: AB_2340593) and 1:500 donkey anti-rat IgG (Jackson Immuno-Research Lab, cat#712-065-153, RRID: AB_2315779). Slides were then incubated with VECTASTAIN Elite ABC HRP Kit (Vector Laboratories, cat#PK-6100) and then developed with ImmPACT DAB Substrate Peroxidase Kit (Vector Laboratories, cat#SK-4105) and counterstained with Hematoxylin QS (Vector Laboratories, cat#H-3404). Slides were scanned at 20x with a Leica Aperio Imager and viewed with Aperio ImageScope software.

Masson’s trichrome stain slides were scanned at 20x with a Leica Aperio Imager and viewed with Aperio ImageScope software. The relative amount of stromal ECM in the endometrium was quantified from these 20x images on Image J’s Color Deconvolution plug-in^127,128^. Four representative integrated density measurements were taken for each mouse and then used to determine each mouse’s average. Fold change in integrated density of HFD to the average of CD was additionally calculated. Picrosirius red stained slides were imaged by hand with brightfield at 20x with a Nikon ECLIPSE N*i* microscope. The relative amount of stromal collagen in the endometrium was quantified from these 20x images using Image J’s Color Deconvolution plug-in. Four representative integrated density measurements were taken for each mouse and then used to determine each mouse’s average. Fold change in integrated density of HFD to the average of CD was additionally calculated. Non-estrous staged uterine H&E-stained slides from HFD and CD *Pten^floxed^;Pax8^Cre/+^* mice were utilized to determine EH disease severity at 10x magnification. Slides were scanned with a Leica Aperio Imager and viewed with Aperio ImageScope software. Total number of glands, percent of normal glands out of total, percent of abnormal glands out of total, average epithelial height from abnormal glands, and average abnormal gland area was calculated per section using Aperio’s ImageScope software. Average epithelial layer height of abnormal glands was calculated by measuring three different places along each of the 20 thickest abnormal glands and then taking each of the 20 glands’ average to then determine the mouse’s average epithelial height (in μm). Average abnormal gland area was determined by measuring the length and height of the 20 largest abnormal glands to obtain the area of the gland (in μm). The 20 glands were then averaged to obtain the mouse’s average gland area.

### Tissue Digestion, Dead Cell and Red Blood Cell (RBC) Removal, and Flow Cytometry Analysis of Macrophages

After 21-30 weeks on CD and HFD, Pten*^fl/+^; Pax8^Cre/+^* estrus-staged mice were euthanized and uteri and blood collected. Uteri were then digested, samples filtered, RBCs lysed, and dead cells removed by MACS as previously described in the above scRNA-seq methods. Alternatively, after cheek bleed blood collection, blood was immediately mixed with 10 U/mL heparin (Sigma-Aldrich, cat#H3393) diluted in RPMI (Gibco, cat#11875-093) until ready for further processing. To then isolate immune cells, blood was likewise incubated with RBC lysis solution. FcR Blocking Reagent (Miltenyi, cat#130-092-575) was added to both the blood and uterus samples before adding the respective antibody combinations. Samples were labelled 1:50 CD45 (Miltenyi, cat#130-123-784, RRID:AB_2802059), 1:50 Propidium Iodide (PI) (BD Biosciences, cat#556463, RRID:AB_2869075), 1:50 F4/80 (BioLegend, cat#123120, RRID:AB_893479), 1:20 CD80 (BioLegend, cat#104713, RRID:AB_313134), 1:40 CD206 (BioLegend, cat#141707, RRID:AB_10896057), as well as isotype controls at a matching concentration including APC Rat IgG2b,τχ (BioLegend, cat#403806, RRID:AB_ 3096358), Alexa Fluor® 488 Rat IgG2a, κ (BioLegend, cat#400525, RRID:AB_2864283), APC Armenian Hamster IgG (BioLegend, cat#400911, RRID:AB_2905474), and APC Rat IgG2a, κ (BioLegend, cat#400511, RRID:AB_2814702). Samples were run on the BD Accuri™ C6 Plus and then files analyzed with FlowJo™ 10.9.0.

### Statistical Analysis

Statistical analyses were performed using GraphPad Prism (Version 10). The *Pten* heterozygous mutant amount of time on diet, mouse age at tissue harvest, uterine weight, uterine-to-body-weight ratio, serum estrone, serum estradiol, serum progesterone, and serum estradiol-to-progesterone ratio, *Pten* control and heterozygous mutant raw and fold change in ECM integrated density from Masson’s trichrome staining and collagen deposition from Picrosirius Red staining, EH disease severity (total number of glands, percent normal and abnormal glands, abnormal gland epithelial height, and abnormal gland area), and *Pten* het flow cytometry samples (uterine and blood CD45^+^, F4/80^+^, PI- dead cells, CD80^+^, CD206^+^, CD80^+^/CD206^+^ ratio) were analyzed with a two-tailed unpaired *t*-test statistic or Welch’s *t-*test statistic when there were unequal variances. Analysis with all six treatment groups of *Pten* control, heterozygous, and homozygous mutant mice or all four *Esr1* control and homozygous mutant mice including age at tissue harvest, time on diet, starting and final mouse weight, uterus weight and uterus-to-body-weight ratio with all treatment groups (Fig. 4 and Supplementary Fig. 1) were calculated using the Brown-Forsythe and Welch ANOVA with Dunnett T3 multiple comparison test statistic. Body weight over time was analyzed by Mixed-effects model with Geisser-Greenhouse correction and Tukey multiple comparison correction test statistic for the six *Pten* treatment groups and multiple unpaired t-test with Holm-Šídák multiple comparison correction for *Pten* heterozygous mutant macrophage flow treatment groups. Glucose tolerance test values were analyzed by a Two-Way ANOVA with Geisser-Greenhouse correction and Tukey multiple comparison correction test statistic. DEGs and differences in cluster abundance were determined by FDR<0.05. Mouse-to-human fibroblast population marker gene analysis was performed using Chi-square statistic with Yates correction and 17,098 mouse-to-human orthologue genes as the total background^129^. The following refers to p-values denoted in figures: *n.s.>0.05; *p<0.05; **p<0.01; ***p<0.001; ****p<0.0001*.

## Data Availability

Sequence data that support the findings of this study have been deposited at the NCBI Gene Expression Omnibus (GSE319670).

## Acknowledgements

We would like to thank the various cores for their services. Thanks to the Van Andel Institute Genomics Core (RRID:SCR_022913), as well as the Bioinformatics and Biostatistics Core (RRID:SCR_024762) for providing single-cell RNA sequencing facilities, services, and analysis. Also, thanks to the Van Andel Institute Pathology and Biorepository Core (RRID:SCR_022912) for their assistance with embedding/sectioning, staining, and scanning our histological samples. Lastly, thanks to Dr. Rachael Sheridan from the Van Andel Institute Flow Cytometry Core for advice on our flow cytometry studies and the Wisconsin National Primate Research Center- Assay Services at the University of Wisconsin-Madison for mass spectrometry serum steroid hormone analysis. We would also like to thank Drs. Tae Hoon Kim and Jae-Wook Jeong at the University of Missouri for providing us with the *Amhr2^Cre^* mouse line.

## Author Contributions

Conceptualization: H.J.S. and R.L.C.; Methodology: H.J.S, K.B., M.W., M.A, and R.L.C.; Investigation: H.J.S., A.Z.B., L.E.W., S.K.H., A.R.A., A.G.L.E., K.L., and R.L.C.; Formal analysis: H.J.S., and K.L.; Data curation: H.J.S., and R.L.C.; Resources: G.W.B., E.N.P., G.H., and J.M.T.; Writing—original draft: H.J.S., K.B., K.L., and R.L.C.; Writing—review and editing: H.J.S., A.Z.B., L.E.W., S.K.H., A.R.A., A.G.L.E., G.W.B., E.N.P., G.H., K.B., M.W., M.A., J.M.T., K.L., and R.L.C.; Funding acquisition: H.J.S and R.L.C.; Supervision: R.L.C.

## Materials & Correspondence

Ronald L. Chandler;

**Supplemental Fig. 1 Experimental timeline and associated metrics of CD and HFD Pten floxed (control), heterozygous, and homozygous mutants**

(a) Experimental timeline for *Pten* floxed (control), heterozygous, and homozygous mutant mice on CD and HFD. This includes both the estrus-staged mice (used for scRNA-sequencing) and well as the rest of the cohort (used for later experiments). (b, c) Mouse age at tissue collection (weeks) and the amount of time on diet (weeks) for the estrus-staged CD and HFD mice that completed sequencing for scRNA-seq as well as for the whole cohort. Sequenced mice included CD (n=2) and HFD (n=2) *Pten ^floxed^* (Cont), CD (n=3) and HFD (n=3) *Pten^fl/+^;Pax8^Cre/+^* (Het), and CD (n=3) and HFD (n=3) *Pten^fl/fl^;Pax8^Cre/+^* (Mut) mice. Total cohort included CD (n=10) and HFD (n=11) *Pten ^floxed^*(Cont), CD (n=15) and HFD (n=16) *Pten^fl/+^;Pax8^Cre/+^* (Het), and CD (n=6) and HFD (n=9) *Pten^fl/fl^;Pax8^Cre/+^* (Mut) mice. Measurement (datapoint) represents each mouse (bar=mean ± s.d.). Brown-Forsythe and Welch ANOVA with Dunnett T3 multiple comparison test statistic. Statistical significance results are only shown for comparisons between diet groups of the same genotype, as well as comparing the highest treatment value to the lowest treatment value, and comparisons that were significant. (d) Average weekly mouse weight (grams) per diet and genotype combination over the experimental time course. Measurement represents a treatment groups’ weekly mean ± s.d. Mixed-effects model with Geisser-Greenhouse correction and Tukey multiple comparison correction test statistic. Statistical significance results are only shown for comparisons between diet groups of the same genotype, as well as comparing the highest CD treatment value to the lowest HFD treatment value (CD Cont to HFD Mut). Moreover, only the initial mouse weights and the first time of statistical significance are displayed on the graph. (e,f) Final mouse body weight for all mice in the cohort for each treatment group with significance displayed by either diet type or genotype. Measurement (datapoint) represents each mouse (bar=mean ± s.d.). Brown-Forsythe and Welch ANOVA with Dunnett T3 multiple comparison test statistic. (g) Glucose tolerance test after 18-24 weeks on diet for all treatment groups. Time point 0 fasting blood glucose (mg/dL) and post-glucose injection blood glucose levels were measured (30, 90, 150, and 210 minutes). Measurement represents a treatment groups’ mean ± s.d. Two-way ANOVA with Geisser-Greenhouse correction and Tukey multiple comparison correction test statistic. Statistical significance results are only shown for comparisons between diet groups of the same genotype, as well as comparing the highest CD treatment value to the lowest HFD treatment value (CD Cont to HFD Mut). (h) Semi-dry uterine weight (mg) at tissue harvest for all mice. Measurement (datapoint) represents each mouse (bar= mean ± s.d.). Brown-Forsythe and Welch ANOVA with Dunnett T3 multiple comparison test statistic. Statistical significance results are only shown for comparisons between diet groups of the same genotype, as well as comparing the highest control genotype (HFD Cont) to the lowest mutant genotype (CD Mut). (i) Semi-dry uterine weight (mg) to mouse final body weight (g) ratio. Measurement (datapoint) represents each mouse (bar= mean ± s.d.). Brown-Forsythe and Welch ANOVA with Dunnett T3 multiple comparison test statistic. Statistical significance results are only shown for comparisons between diet groups of the same genotype as well as comparisons that were significant. *\*p<0.05; **p<0.01; ***p<0.001; ****p<0.0001*

**Supplemental Fig. 2 scRNAseq-identified doublets in isolated endometrial samples**

(a) Stacked bar graph representing the percent of cells in each sample that were identified as doublets using scDblFinder. (b) UMAP of where the doublets were identified in the endometrial sample clusters of all six treatment groups. Clusters with high doublet scores are listed as “#.D”. These clusters (11.D, 13.D, and 19.D) were removed from further analysis. (c) Violin plot displaying the doublet score composition for each identified endometrial cluster.

**Supplemental Fig. 3 Cluster abundance and differential gene expression between all treatment groups**

(a) UMAP illustrations of endometrial cell type clusters over all scRNA-seq conditions: CD *Pten^floxed^*control (n=2), HFD *Pten^floxed^* control (n=2), CD *Pten^fl/+^; Pax8^Cre/+^* het (n=3), HFD *Pten^fl/+^; Pax8^Cre/+^*het (n=3), CD *Pten^fl/fl^; Pax8^Cre/+^* mut (n=3), and HFD *Pten^fl/fl^; Pax8^Cre/+^*mut (n=3). (b) Difference in cluster abundance between *Pten^fl/fl^; Pax8^Cre/+^* and *Pten^floxed^* control mice on either CD or HFD. Difference measured in fold change (FC) with significance of FDR<0.05. (b) Number of DEGs for each cell type in HFD versus CD *Pten* heterozygous mutant mice. (d) Number of DEGs for each cell type in HFD versus CD *Pten* homozygous mutant mice. (e) Number of DEGs for each cell type in *Pten* heterozygous mutant versus *Pten* floxed control mice on CD. (f) Number of DEGs for each cell type in *Pten* homozygous mutant versus *Pten* floxed control mice on CD. (g) Number of DEGs for each cell type in *Pten* heterozygous mutant versus *Pten* floxed control mice on HFD. (h) Number of DEGs for each cell type in *Pten* homozygous mutant versus *Pten* floxed control mice on HFD. DEG significance set at FDR<0.05.

**Supplemental Fig. 4 Gene expression profiles for the three major fibroblast subtypes**

(a) Euler plot showing overlap of top 50 marker genes in fibroblast Cluster 4 (Fibroblast I), Cluster 7 and Cluster 9 (Fibroblast II and III) in CD- or HFD-treated control conditions as determined by CyteType^123^. (b) Euler plot for marker gene comparison between our mouse Cluster 4 (Fibroblast I) population to the human “C7 Fibroblast” population^44^. Overlap of mouse-to-human marker genes was performed using Chi-square statistic with Yates correction and 17,098 mouse-to-human orthologue genes as the total background^129^. (c) Bubbleplot of selected marker gene expression in Gsn^+^ Fibroblast I (Cluster 4) and Aldh1a2^+^ Fibroblast II/III (Clusters 7/9). (d) Heatmap of top 200 genes changing across the three fibroblast clusters from a stromal pseudotime analysis. (e) Heatmap from transcription factor (TF) activity analysis for fibroblast Clusters 4 (Gsn^+^ Fibroblast), 7 and 9 (Aldh1a2^+^ Fibroblast II/III) for both CD and HFD conditions. Significance was tested by comparing Cluster 4 (Gsn^+^ Fibroblast) to either Cluster 7 or Cluster 9 (Aldh1a2^+^ Fibroblast II or III) for each diet. All TFs with an FDR ≤ 0.05 for at least 2 comparisons were plotted as heatmaps. *\*FDR<0.05; **FDR<0.01; ***FDR<0.001; ****FDR<0.0001*

**Supplemental Fig. 5 Additional study metrics for ERα cKO mice**

(a) Mouse weight (g) of *Esr1^fl/fl^; Amhr2^+/+^*(cont) and *Esr1^fl/fl^; Amhr2^Cre/+^* (mut) mice at the start of the experiment for both CD- and HFD-treated conditions. Measurement (datapoint) represents each mouse (bar=mean ± s.d.). Brown-Forsythe and Welch ANOVA with Dunnett T3 multiple comparison test statistic. (b,c) Amount of time on diet (weeks) and mouse age at tissue collection (weeks) for the CD and HFD-treated control mice. CD (n=9) and HFD (n=7) estrus-staged control, and CD (n=9) and HFD (n=9) diestrus-staged mutant mice. Measurement (datapoint) represents each mouse (bar=mean ± s.d.). Brown-Forsythe and Welch ANOVA with Dunnett T3 multiple comparison test statistic. *\*p<0.05; **p<0.01; ***p<0.001; ****p<0.0001*

**Supplemental Fig. 6 Cluster 5 subclustering for monocytes, macrophages and dendritic cells**

(a) UMAP plot showing the subclusters of Cluster 5, where macrophages were identified in subclusters 5.2 and 5.5. Subclustering performed using all treatment groups. (b) Heatmap of immune cell marker genes confirming cluster 5 cell type annotations from HFD and CD *Pten* floxed control samples.

**Supplemental Fig. 7 *Mki67* expression and cell cycle stage in epithelial cells across conditions**

(a) Stacked bar graph of percent of Cluster 2 that is Epithelium VII versus Epithelium I in CD- and HFD-treated controls. (b) Stacked bar graph of percent of total Epithelium VII within the different clusters. (c) Heatmap of epithelial stem cell and proliferation (*Mki67*) marker expression in the Epithelium VII versus the other six epithelium across all treatment groups. (d) Stacked bar graph depicting percent of stem cell-like Epithelium VII out of total epithelium in CD-treated and HFD-treated controls. (e-h) UMAPs and graphs of the six epithelial clusters with expression levels of *Mki67* or percent of cells in each epithelial cluster and/or treatment group that express *Mki67*. (i-k) Stacked bar graphs of percent or number of cells in each cell cycle stage for each epithelial cluster and/or treatment group. Note: CD-treated controls only had one cell in Cluster 18.

**Supplemental Fig. 8 CD and HFD *Pten* heterozygous mutant cohort descriptions and additional macrophage flow cytometry data**

(a,b) Amount of time on diet (weeks) and age at tissue collection (weeks) for the non-estrous staged CD (n=10) and HFD (n=12) *Pten* heterozygous mutant mice, which were in the same cohort as the estrus-staged heterozygous mice selected for scRNA-seq (see Supplementary Fig. 1). Measurement (datapoint) represents each mouse (bar=mean ± s.d.). Two-tailed unpaired *t*-test statistic. (c) Semi-dry uterine weight (mg) at tissue harvest. Measurement (datapoint) represents each mouse (bar= mean ± s.d.). Two-tailed unpaired *t*-test statistic with Welch’s correction. (d) Semi-dry uterine weight (mg) to mouse final body weight (g) ratio. Measurement (datapoint) represents each mouse (bar= mean ± s.d.). Two-tailed unpaired *t*-test statistic with Welch’s correction. (e-h) Serum estrone (CD n=8; HFD n=6), estradiol (CD n=6; HFD n=4), progesterone (CD n=9; HFD n=7), and estradiol-to-progesterone ratio (CD n=6; HFD n=4) levels were measured (ng/ml) in estrus-staged *Pten* heterozygous mutant mice. Some samples had undetectable levels of estrone and estradiol. Measurement (datapoint) represents each mouse (bar= mean ± s.d.). Two-tailed unpaired *t*-test statistic with Welch’s correction as needed. (i) Experimental timeline for CD and HFD-treated *Pten* heterozygous mutant mice used for macrophage flow cytometry. (j) Average weekly mouse weight (grams) for both CD- (n=8) and HFD- (n=9) treated mice over the experimental time course. Measurement represents a diet’s weekly mean ± s.d. Multiple unpaired *t*-test statistic with Holm-Šídák multiple comparison correction. After reaching statistical significance, asterisks were only added when an increase in significance occurred (all weeks were singificant after 2 weeks on diet). (k) Glucose tolerance test after 17.5-22 weeks on diet (with exception of one outlier tested at 26 weeks). Time point 0 fasting blood glucose (mg/dL) and post-glucose injection blood glucose levels were measured (30, 90, 150, and 210 minutes). Measurement represents a diet’s mean ± s.d. Two-way ANOVA with Geisser-Greenhouse correction and Tukey multiple comparison correction test statistic. (l) Average mouse body weight (grams) at the time of tissue harvest for the estrus-staged CD (n=5) and HFD (n=8) *Pten* heterozygous mutant mice. Measurement (datapoint) represents each mouse (bar=mean ± s.d.). Two-tailed unpaired *t*-test statistic. (m,n) Amount of time on diet (weeks) and age at tissue collection (weeks). Measurement (datapoint) represents each mouse (bar=mean ± s.d.). Two-tailed unpaired *t*-test statistic. (o,p) Representative uterine and blood flow cytometry gating scheme for F4/80^+^ macrophages. Analysis performed with FlowJo™ 10.9.0. (q) Illustration of how F4/80^+^ monocytes exit the blood vessel and enter the uterus where they differentiate into F4/80^+^ macrophages. We further utilized CD80 as a marker for pro-inflammatory macrophages and CD206 for anti-inflammatory macrophages. Graphic created with BioRender. (r-w) Percent (%) of CD80^+^ or CD206^+^ cells from all F4/80^+^ live single cells in the uterus or the blood and subsequent ratio of CD80^+^ to CD206^+^ cells (HFD n=7 for CD80^+^ and CD206^+^; HFD n=6 for ratio). Measurement (datapoint) represents each mouse (bar=mean ± s.d.). Two-tailed unpaired *t*-test statistic with Welch’s correction as needed. *\*p<0.05; **p<0.01; ***p<0.001; ****p<0.0001*

**Supplemental Fig. 9 Normal epithelium morphology persists, as previously observed in HFD-treated endometrium**

(a,b) Representative H&E stain of mouse endometrium for CD and HFD *Pten* floxed controls. Main image= 20x magnification; inset=40x magnification; scalebar=200 μm. Arrows indicate endometrial glands. (c,d) Representative KRT8 IHC epithelial stain of mouse endometrium for CD and HFD controls. Main image= 20x magnification; inset=40x magnification; scalebar=200 μm. (e,f) Representative PTEN stain of mouse endometrium for CD and HFD controls. Main image= 20x magnification; inset=40x magnification; scalebar=200 μm. Arrows indicate PTEN positive-staining endometrial glands.

## References

1. Mazur E, Large M, Demayo F. Chapter 24. Human Oviduct and Endometrium. In: Knobil and Neill’s Physiology of Reproduction. Fourth Edition. 2015. doi:10.1016/B978-0-12-397175-3.00024-7

2. Flynn L, Byrne B, Carton J, Kelehan P, O’Herlihy C, O’Farrelly C. Menstrual cycle dependent fluctuations in NK and T-lymphocyte subsets from non-pregnant human endometrium. Am J Reprod Immunol. 2000;43(4):209–217. doi:10.1111/j.8755-8920.2000.430405.x

3. Agostinis C, Mangogna A, Bossi F, Ricci G, Kishore U, Bulla R. Uterine Immunity and Microbiota: A Shifting Paradigm. Front Immunol. 2019;10:2387. doi:10.3389/fimmu.2019.02387

4. Tatematsu BK, Sojka DK. Tissue-resident natural killer cells derived from conventional natural killer cells are regulated by progesterone in the uterus. Mucosal Immunology. 2025;18(2):390–401. doi:10.1016/j.mucimm.2024.12.009

5. Cousins FL, Kirkwood PM, Saunders PTK, Gibson DA. Evidence for a dynamic role for mononuclear phagocytes during endometrial repair and remodelling. Sci Rep. 2016;6(1):36748. doi:10.1038/srep36748

6. Cooke PS, Buchanan DL, Young P, et al. Stromal estrogen receptors mediate mitogenic effects of estradiol on uterine epithelium. Proc Natl Acad Sci U S A. 1997;94(12):6535–6540. doi:10.1073/pnas.94.12.6535

7. Winuthayanon W, Lierz SL, Delarosa KC, et al. Juxtacrine Activity of Estrogen Receptor α in Uterine Stromal Cells is Necessary for Estrogen-Induced Epithelial Cell Proliferation. Sci Rep. 2017;7(1):8377. doi:10.1038/s41598-017-07728-1

8. Winuthayanon W, Hewitt SC, Orvis GD, Behringer RR, Korach KS. Uterine epithelial estrogen receptor α is dispensable for proliferation but essential for complete biological and biochemical responses. Proceedings of the National Academy of Sciences. 2010;107(45):19272–19277. doi:10.1073/pnas.1013226107

9. Binder AK, Winuthayanon W, Hewitt SC, Couse JF, Korach KS. Chapter 25 Steroid Receptors in the Uterus and Ovary. In: Knobil and Neill’s Physiology of Reproduction. Fourth Edition. 2015:1099–1193. doi:10.1016/B978-0-12-397175-3.00025-9

10. Tong W, Pollard JW. Progesterone inhibits estrogen-induced cyclin D1 and cdk4 nuclear translocation, cyclin E- and cyclin A-cdk2 kinase activation, and cell proliferation in uterine epithelial cells in mice. Mol Cell Biol. 1999;19(3):2251–2264. doi:10.1128/MCB.19.3.2251

11. Martin L, Finn CA. HORMONAL REGULATION OF CELL DIVISION IN EPITHELIAL AND CONNECTIVE TISSUES OF THE MOUSE UTERUS. Published online July 1, 1968. doi:10.1677/joe.0.0410363

12. Kurita T, Young P, Brody JR, Lydon JP, O’Malley BW, Cunha GR. Stromal progesterone receptors mediate the inhibitory effects of progesterone on estrogen-induced uterine epithelial cell deoxyribonucleic acid synthesis. Endocrinology. 1998;139(11):4708–4713. doi:10.1210/endo.139.11.6317

13. Franco HL, Rubel CA, Large MJ, et al. Epithelial progesterone receptor exhibits pleiotropic roles in uterine development and function. The FASEB Journal. 2012;26(3):1218–1227. doi:10.1096/fj.11-193334

14. Kurita T, Lee K jun, Cooke PS, Taylor JA, Lubahn DB, Cunha GR. Paracrine Regulation of Epithelial Progesterone Receptor by Estradiol in the Mouse Female Reproductive Tract1. Biology of Reproduction. 2000;62(4):821–830. doi:10.1093/biolreprod/62.4.821

15. Wira CR, Fahey JV, Sentman CL, Pioli PA, Shen L. Innate and adaptive immunity in female genital tract: cellular responses and interactions. Immunological Reviews. 2005;206(1):306–335. doi:10.1111/j.0105-2896.2005.00287.x

16. MacLean JA, Hayashi K. Progesterone Actions and Resistance in Gynecological Disorders. Cells. 2022;11(4):647. doi:10.3390/cells11040647

17. Pawar S, Hantak AM, Bagchi IC, Bagchi MK. Minireview: Steroid-Regulated Paracrine Mechanisms Controlling Implantation. Mol Endocrinol. 2014;28(9):1408–1422. doi:10.1210/me.2014-1074

18. Ferenczy A, Gelfand M. The biologic significance of cytologic atypia in progestogen-treated endometrial hyperplasia. American Journal of Obstetrics and Gynecology. 1989;160(1):126–131. doi:10.1016/0002-9378(89)90103-8

19. Gunderson CC, Fader AN, Carson KA, Bristow RE. Oncologic and reproductive outcomes with progestin therapy in women with endometrial hyperplasia and grade 1 adenocarcinoma: a systematic review. Gynecol Oncol. 2012;125(2):477–482. doi:10.1016/j.ygyno.2012.01.003

20. Deligdisch L. Hormonal Pathology of the Endometrium. Modern Pathology. 2000;13(3):285–294. doi:10.1038/modpathol.3880050

21. Grodstein F, Goldman MB, Cramer DW. Body mass index and ovulatory infertility. Epidemiology. 1994;5(2):247–250. doi:10.1097/00001648-199403000-00016

22. Green BB, Weiss NS, Daling JR. Risk of ovulatory infertility in relation to body weight*. Fertility and Sterility. 1988;50(5):721–726. doi:10.1016/S0015-0282(16)60305-9

23. Reed BG, Carr BR. The Normal Menstrual Cycle and the Control of Ovulation. In: Feingold KR, Ahmed SF, Anawalt B, et al., eds. Endotext. MDText.com, Inc.; 2000. Accessed July 11, 2025. http://www.ncbi.nlm.nih.gov/books/NBK279054/

24. Quennell JH, Mulligan AC, Tups A, et al. Leptin Indirectly Regulates Gonadotropin-Releasing Hormone Neuronal Function. Endocrinology. 2009;150(6):2805–2812. doi:10.1210/en.2008-1693

25. Quennell JH, Howell CS, Roa J, Augustine RA, Grattan DR, Anderson GM. Leptin Deficiency and Diet-Induced Obesity Reduce Hypothalamic Kisspeptin Expression in Mice. Endocrinology. 2011;152(4):1541–1550. doi:10.1210/en.2010-1100

26. Jain A, Polotsky AJ, Rochester D, et al. Pulsatile Luteinizing Hormone Amplitude and Progesterone Metabolite Excretion Are Reduced in Obese Women. The Journal of Clinical Endocrinology & Metabolism. 2007;92(7):2468–2473. doi:10.1210/jc.2006-2274

27. Santoro N, Lasley B, McConnell D, et al. Body Size and Ethnicity Are Associated with Menstrual Cycle Alterations in Women in the Early Menopausal Transition: The Study of Women’s Health across the Nation (SWAN) Daily Hormone Study. The Journal of Clinical Endocrinology & Metabolism. 2004;89(6):2622–2631. doi:10.1210/jc.2003-031578

28. Skalski HJ, Arendt AR, Harkins SK, et al. Key Considerations for Studying the Effects of High-Fat Diet on the Nulligravid Mouse Endometrium. J Endocr Soc. 2024;8(7):bvae104. doi:10.1210/jendso/bvae104

29. Wilson MR, Skalski H, Reske JJ, et al. Obesity alters the mouse endometrial transcriptome in a cell context-dependent manner. Reprod Biol Endocrinol. 2022;20(1):163. doi:10.1186/s12958-022-01030-0

30. Maxwell GL, Risinger JI, Gumbs C, et al. Mutation of the PTEN tumor suppressor gene in endometrial hyperplasias. Cancer Res. 1998;58(12):2500–2503.

31. Konopka B, Paszko Z, Janiec-Jankowska A, Goluda M. Assessment of the quality and frequency of mutations occurrence in PTEN gene in endometrial carcinomas and hyperplasias. Cancer Letters. 2002;178(1):43–51. doi:10.1016/S0304-3835(01)00815-1

32. Lac V, Nazeran TM, Tessier-Cloutier B, et al. Oncogenic mutations in histologically normal endometrium: the new normal? The Journal of Pathology. 2019;249(2):173–181. doi:10.1002/path.5314

33. Lee H, Choi HJ, Kang CS, Lee HJ, Lee WS, Park CS. Expression of miRNAs and PTEN in endometrial specimens ranging from histologically normal to hyperplasia and endometrial adenocarcinoma. Modern Pathology. 2012;25(11):1508–1515. doi:10.1038/modpathol.2012.111

34. Russo A, Czarnecki AA, Dean M, et al. PTEN loss in the fallopian tube induces hyperplasia and ovarian tumor formation. Oncogene. 2018;37(15):1976–1990. doi:10.1038/s41388-017-0097-8

35. Fei L, Chen H, Ma L, et al. Systematic identification of cell-fate regulatory programs using a single-cell atlas of mouse development. Nat Genet. 2022;54(7):1051–1061. doi:10.1038/s41588-022-01118-8

36. Zhang L, Long W, Xu W, Chen X, Zhao X, Wu B. Digital Cell Atlas of Mouse Uterus: From Regenerative Stage to Maturational Stage. Front Genet. 2022;13. doi:10.3389/fgene.2022.847646

37. Garcia-Flores V, Romero R, Peyvandipour A, et al. A single-cell atlas of murine reproductive tissues during preterm labor. Cell Rep. 2023;42(1):111846. doi:10.1016/j.celrep.2022.111846

38. Winkler I, Tolkachov A, Lammers F, et al. The cycling and aging mouse female reproductive tract at single-cell resolution. Cell. 2024;187(4):981–998.e25. doi:10.1016/j.cell.2024.01.021

39. Kaeding AJ, Barwe SP, Gopalakrishnapillai A, et al. Mesothelin is a novel cell surface disease marker and potential therapeutic target in acute myeloid leukemia. Blood Advances. 2021;5(9):2350–2361. doi:10.1182/bloodadvances.2021004424

40. Hu C, Li T, Xu Y, et al. CellMarker 2.0: an updated database of manually curated cell markers in human/mouse and web tools based on scRNA-seq data. Nucleic Acids Research. 2023;51(D1):D870–D876. doi:10.1093/nar/gkac947

41. Blaisdell A, Crequer A, Columbus D, et al. Neutrophils Oppose Uterine Epithelial Carcinogenesis via Debridement of Hypoxic Tumor Cells. Cancer Cell. 2015;28(6):785–799. doi:10.1016/j.ccell.2015.11.005

42. Hinz B, Gabbiani G. Mechanisms of force generation and transmission by myofibroblasts. Current Opinion in Biotechnology. 2003;14(5):538–546. doi:10.1016/j.copbio.2003.08.006

43. Arora PD, McCulloch CAG. Dependence of collagen remodelling on α-smooth muscle actin expression by fibroblasts. Journal of Cellular Physiology. 1994;159(1):161–175. doi:10.1002/jcp.1041590120

44. Ulrich ND, Vargo A, Ma Q, et al. Cellular heterogeneity and dynamics of the human uterus in healthy premenopausal women. Proceedings of the National Academy of Sciences. 2024;121(45):e2404775121. doi:10.1073/pnas.2404775121

45. Wang SSY, Tang H, Loe MWC, Yeo SC, Javaid MM. Complements and Their Role in Systemic Disorders. Cureus. 16(1):e52991. doi:10.7759/cureus.52991

46. Lian S, Liu J, Yang Y, Zhu G, Xia P. The life-and-death struggle between the complement system and pathogens: Mechanisms of elimination, evasion tactics, and translational potential. Virulence. 16(1):2553781. doi:10.1080/21505594.2025.2553781

47. Li Y, Zhao J, Yin Y, Li K, Zhang C, Zheng Y. The Role of IL-6 in Fibrotic Diseases: Molecular and Cellular Mechanisms. Int J Biol Sci. 2022;18(14):5405–5414. doi:10.7150/ijbs.75876

48. Wilkens J, Male V, Ghazal P, et al. Uterine Natural Killer cells regulate endometrial bleeding in women and are suppressed by the progesterone receptor modulator asoprisnil. J Immunol. 2013;191(5):10.4049/jimmunol.1300958. doi:10.4049/jimmunol.1300958

49. Wernig G, Chen SY, Cui L, et al. Unifying mechanism for different fibrotic diseases. Proceedings of the National Academy of Sciences. 2017;114(18):4757–4762. doi:10.1073/pnas.1621375114

50. Yang M, Jiang C, Li L, Xing H, Hong L. Expression of CD47 in Endometrial Cancer and Its Clinicopathological Significance. Journal of Oncology. 2022;2022(1):7188972. doi:10.1155/2022/7188972

51. DeMayo FJ, Lydon JP. 90 YEARS OF PROGESTERONE: New insights into progesterone receptor signaling in the endometrium required for embryo implantation. Published online July 1, 2020. doi:10.1530/JME-19-0212

52. Okada H, Nakajima T, Sanezumi M, Ikuta A, Yasuda K, Kanzaki H. Progesterone enhances interleukin-15 production in human endometrial stromal cells in vitro. J Clin Endocrinol Metab. 2000;85(12):4765–4770. doi:10.1210/jcem.85.12.7023

53. Wilkens J, Male V, Ghazal P, et al. Uterine Natural Killer cells regulate endometrial bleeding in women and are suppressed by the progesterone receptor modulator asoprisnil. J Immunol. 2013;191(5):10.4049/jimmunol.1300958. doi:10.4049/jimmunol.1300958

54. Li Q, Kannan A, DeMayo FJ, et al. The Antiproliferative Action of Progesterone in Uterine Epithelium Is Mediated by Hand2. Science. 2011;331(6019):912–916. doi:10.1126/science.1197454

55. Wang Y, Chen Z, Wang R, Song A, Zhang Y. Obesity and oxidative stress: potential mechanisms in endometrial disorders. Front Endocrinol (Lausanne). 17:1709556. doi:10.3389/fendo.2026.1709556

56. Joo E, Yamada KM. Cell Adhesion and Movement. In: Stem Cell Biology and Tissue Engineering in Dental Sciences. Academic Press; 2015:61–72. doi:10.1016/B978-0-12-397157-9.00005-9

57. Gao D, Hu S, Zheng X, et al. SOD3 Is Secreted by Adipocytes and Mitigates High-Fat Diet-Induced Obesity, Inflammation, and Insulin Resistance. Antioxid Redox Signal. 2020;32(3):193–212. doi:10.1089/ars.2018.7628

58. Zhang L, Han C, Ye F, et al. Plasma Gelsolin Induced Glomerular Fibrosis via the TGF-β1/Smads Signal Transduction Pathway in IgA Nephropathy. Int J Mol Sci. 2017;18(2):390. doi:10.3390/ijms18020390

59. Wynn TA, Ramalingam TR. Mechanisms of fibrosis: therapeutic translation for fibrotic disease. Nat Med. 2012;18(7):1028–1040. doi:10.1038/nm.2807

60. Bernau K, Leet JP, Bruhn EM, Tubbs AJ, Zhu T, Sandbo N. Expression of serum response factor in the lung mesenchyme is essential for development of pulmonary fibrosis. Am J Physiol Lung Cell Mol Physiol. 2021;321(1):L174–L188. doi:10.1152/ajplung.00323.2020

61. Matsuzaki S, Pouly JL, Canis M. Persistent activation of signal transducer and activator of transcription 3 via interleukin-6 trans-signaling is involved in fibrosis of endometriosis. Hum Reprod. 2022;37(7):1489–1504. doi:10.1093/humrep/deac098

62. Liu L, Chen G, Chen T, et al. si-SNHG5-FOXF2 inhibits TGF-β1-induced fibrosis in human primary endometrial stromal cells by the Wnt/β-catenin signalling pathway. Stem Cell Res Ther. 2020;11:479. doi:10.1186/s13287-020-01990-3

63. Kriseman M, Monsivais D, Agno J, Masand RP, Creighton CJ, Matzuk MM. Uterine double-conditional inactivation of Smad2 and Smad3 in mice causes endometrial dysregulation, infertility, and uterine cancer. Proceedings of the National Academy of Sciences. 2019;116(9):3873–3882. doi:10.1073/pnas.1806862116

64. Simmen RCM, Heard ME, Simmen AM, et al. The Krüppel-Like Factors in Female Reproductive System Pathologies. J Mol Endocrinol. 2015;54(2):R89–R101. doi:10.1530/JME-14-0310

65. Gao Y, Li S, Li Q. Uterine epithelial cell proliferation and endometrial hyperplasia: evidence from a mouse model. Mol Hum Reprod. 2014;20(8):776–786. doi:10.1093/molehr/gau033

66. Wynn TA, Ramalingam TR. Mechanisms of fibrosis: therapeutic translation for fibrotic disease. Nat Med. 2012;18(7):1028–1040. doi:10.1038/nm.2807

67. Jamin SP, Arango NA, Mishina Y, Hanks MC, Behringer RR. Requirement of Bmpr1a for Müllerian duct regression during male sexual development. Nat Genet. 2002;32(3):408–410. doi:10.1038/ng1003

68. Hewitt SC, Kissling GE, Fieselman KE, Jayes FL, Gerrish KE, Korach KS. Biological and biochemical consequences of global deletion of exon 3 from the ERα gene. FASEB J. 2010;24(12):4660–4667. doi:10.1096/fj.10-163428

69. Porteous R, Herbison AE. Genetic Deletion of Esr1 in the Mouse Preoptic Area Disrupts the LH Surge and Estrous Cyclicity. Endocrinology. 2019;160(8):1821–1829. doi:10.1210/en.2019-00284

70. Dupont S, Krust A, Gansmuller A, Dierich A, Chambon P, Mark M. Effect of single and compound knockouts of estrogen receptors α (ERα) and β (ERβ) on mouse reproductive phenotypes. Development. 2000;127(19):4277–4291. doi:10.1242/dev.127.19.4277

71. Laws MJ, Kannan A, Pawar S, Haschek WM, Bagchi MK, Bagchi IC. Dysregulated estrogen receptor signaling in the hypothalamic-pituitary-ovarian axis leads to ovarian epithelial tumorigenesis in mice. PLoS Genet. 2014;10(3):e1004230. doi:10.1371/journal.pgen.1004230

72. Winuthayanon W, Lierz SL, Delarosa KC, et al. Juxtacrine Activity of Estrogen Receptor α in Uterine Stromal Cells is Necessary for Estrogen-Induced Epithelial Cell Proliferation. Sci Rep. 2017;7:8377. doi:10.1038/s41598-017-07728-1

73. Davidson S, Coles M, Thomas T, et al. Fibroblasts as immune regulators in infection, inflammation and cancer. Nat Rev Immunol. 2021;21(11):704–717. doi:10.1038/s41577-021-00540-z

74. Pittet MJ, Nahrendorf M, Swirski FK. The journey from stem cell to macrophage. Ann N Y Acad Sci. 2014;1319(1):1–18. doi:10.1111/nyas.12393

75. Liu D, Wang J, Zhao G, et al. CSF1-associated decrease in endometrial macrophages may contribute to Asherman’s syndrome. American Journal of Reproductive Immunology. 2020;83(1):e13191. doi:10.1111/aji.13191

76. Fu B, Li X, Sun R, et al. Natural killer cells promote immune tolerance by regulating inflammatory TH17 cells at the human maternal–fetal interface. Proceedings of the National Academy of Sciences. 2013;110(3):E231–E240. doi:10.1073/pnas.1206322110

77. Hanna J, Goldman-Wohl D, Hamani Y, et al. Decidual NK cells regulate key developmental processes at the human fetal-maternal interface. Nat Med. 2006;12(9):1065–1074. doi:10.1038/nm1452

78. Cai SF, Fehniger TA, Dai X, Cao X, Ley TJ. Differential Regulation of Granzyme B and C Expression in Murine Cytotoxic T and NK Cells. Blood. 2007;110(11):2291. doi:10.1182/blood.V110.11.2291.2291

79. Bourayou E, Golub R. Inflammatory-driven NK cell maturation and its impact on pathology. Front Immunol. 2022;13:1061959. doi:10.3389/fimmu.2022.1061959

80. Gorsek Sparovec T, Markert UR, Reif P, et al. The fate of human SUSD2+ endometrial mesenchymal stem cells during decidualization. Stem Cell Res. 2022;60:102671. doi:10.1016/j.scr.2022.102671

81. Cousins FL, Filby CE, Gargett CE. Endometrial Stem/Progenitor Cells–Their Role in Endometrial Repair and Regeneration. Front Reprod Health. 2022;3:811537. doi:10.3389/frph.2021.811537

82. Seishima R, Leung C, Yada S, et al. Neonatal Wnt-dependent Lgr5 positive stem cells are essential for uterine gland development. Nat Commun. 2019;10:5378. doi:10.1038/s41467-019-13363-3

83. Zheng SC, Stein-O’Brien G, Augustin JJ, et al. Universal prediction of cell-cycle position using transfer learning. Genome Biol. 2022;23(1):41. doi:10.1186/s13059-021-02581-y

84. Bharti R, Dey G, Lin F, Lathia J, Reizes O. CD55 in cancer: Complementing functions in a non-canonical manner. Cancer Letters. 2022;551:215935. doi:10.1016/j.canlet.2022.215935

85. Bellone S, Roque D, Cocco E, et al. Downregulation of membrane complement inhibitors CD55 and CD59 by siRNA sensitises uterine serous carcinoma overexpressing Her2/neu to complement and antibody-dependent cell cytotoxicity in vitro: implications for trastuzumab-based immunotherapy. Br J Cancer. 2012;106(9):1543–1550. doi:10.1038/bjc.2012.132

86. Poon CE, Madawala RJ, Dowland SN, Murphy CR. Nectin-3 Is Increased in the Cell Junctions of the Uterine Epithelium at Implantation. Reprod Sci. 2016;23(11):1580–1592. doi:10.1177/1933719116648216

87. Luo L, Zhang W, You S, et al. The role of epithelial cells in fibrosis: Mechanisms and treatment. Pharmacological Research. 2024;202:107144. doi:10.1016/j.phrs.2024.107144

88. Austyn JM, Gordon S. F4/80, a monoclonal antibody directed specifically against the mouse macrophage. Eur J Immunol. 1981;11(10):805–815. doi:10.1002/eji.1830111013

89. Singh G, Cue L, Puckett Y. Endometrial Hyperplasia. In: StatPearls. StatPearls Publishing; 2025. Accessed July 15, 2025. http://www.ncbi.nlm.nih.gov/books/NBK560693/

90. Giuliani E, Schon SB, Yang K, et al. Obesity-induced follicular phase endometrial proteome dysregulation in a well-phenotyped population. F S Sci. 2022;3(4):367–375. doi:10.1016/j.xfss.2022.06.002

91. Kirkwood PM, Gibson DA, Smith JR, et al. Single-cell RNA sequencing redefines the mesenchymal cell landscape of mouse endometrium. FASEB J. 2021;35(4):e21285. doi:10.1096/fj.202002123R

92. Garcia-Alonso L, Handfield LF, Roberts K, et al. Mapping the temporal and spatial dynamics of the human endometrium in vivo and in vitro. Nat Genet. 2021;53(12):1698–1711. doi:10.1038/s41588-021-00972-2

93. Garcia-Alonso L, Handfield LF, Roberts K, et al. Mapping the temporal and spatial dynamics of the human endometrium in vivo and in vitro. Nat Genet. 2021;53(12):1698–1711. doi:10.1038/s41588-021-00972-2

94. Marečková M, Garcia-Alonso L, Moullet M, et al. An integrated single-cell reference atlas of the human endometrium. Nat Genet. 2024;56(9):1925–1937. doi:10.1038/s41588-024-01873-w

95. Sahin Ersoy G, Zolbin MM, Cosar E, Moridi I, Mamillapalli R, Taylor HS. CXCL12 Promotes Stem Cell Recruitment and Uterine Repair after Injury in Asherman’s Syndrome. Mol Ther Methods Clin Dev. 2017;4:169–177. doi:10.1016/j.omtm.2017.01.001

96. Okada H, Tsuzuki T, Murata H. Decidualization of the human endometrium. Reprod Med Biol. 2018;17(3):220–227. doi:10.1002/rmb2.12088

97. Laurila JP, Laatikainen LE, Castellone MD, Laukkanen MO. SOD3 reduces inflammatory cell migration by regulating adhesion molecule and cytokine expression. PLoS One. 2009;4(6):e5786. doi:10.1371/journal.pone.0005786

98. Sundstrom SA, Komm BS, Ponce-de-Leon H, Yi Z, Teuscher C, Lyttle CR. Estrogen regulation of tissue-specific expression of complement C3. J Biol Chem. 1989;264(28):16941–16947.

99. Brown EO, Sundstrom SA, Komm BS, Yi Z, Teuscher C, Richard Lyttle C. Progesterone Regulation of Estradiol-Induced Rat Uterine Secretory Protein, Complement C31. Biol Reprod. 1990;42(4):713–719. doi:10.1095/biolreprod42.4.713

100. Scott NA, Mohiyiddeen L, Mariano LL, et al. Macrophages in the uterus are functionally specialised and continually replenished from the circulation. bioRxiv. Preprint posted online September 3, 2021:2021.09.02.458731. doi:10.1101/2021.09.02.458731

101. Yang Z, Kong B, Mosser DM, Zhang X. TLRs, macrophages, and NK cells: Our understandings of their functions in uterus and ovary. International Immunopharmacology. 2011;11(10):1442–1450. doi:10.1016/j.intimp.2011.04.024

102. Hartmann N, Giese NA, Giese T, et al. Prevailing Role of Contact Guidance in Intrastromal T-cell Trapping in Human Pancreatic Cancer. Clin Cancer Res. 2014;20(13):3422–3433. doi:10.1158/1078-0432.CCR-13-2972

103. Epplein M, Reed SD, Voigt LF, Newton KM, Holt VL, Weiss NS. Risk of Complex and Atypical Endometrial Hyperplasia in Relation to Anthropometric Measures and Reproductive History. Am J Epidemiol. 2008;168(6):563–570. doi:10.1093/aje/kwn168

104. Sanderson PA, Critchley HOD, Williams ARW, Arends MJ, Saunders PTK. New concepts for an old problem: the diagnosis of endometrial hyperplasia. Human Reproduction Update. 2017;23(2):232–254. doi:10.1093/humupd/dmw042

105. Vijayaraghavan A, Jadhav C, Pradeep B, Bindu H, Kumaran S. A Histopathological Study of Endometrial Biopsy Samples in Abnormal Uterine Bleeding. Cureus. 14(11):e31264. doi:10.7759/cureus.31264

106. Prior JC. Perimenopause: The Complex Endocrinology of the Menopausal Transition. Accessed February 4, 2026. 10.1210/edrv.19.4.0341

107. Lesche R, Groszer M, Gao J, et al. Cre/loxP-mediated inactivation of the murine Pten tumor suppressor gene. Genesis. 2002;32(2):148–149. doi:10.1002/gene.10036

108. Bouchard M, Souabni A, Mandler M, Neubüser A, Busslinger M. Nephric lineage specification by Pax2 and Pax8. Genes Dev. 2002;16(22):2958–2970. doi:10.1101/gad.240102

109. Amezquita RA, Lun ATL, Becht E, et al. Orchestrating single-cell analysis with Bioconductor. Nat Methods. 2020;17(2):137–145. doi:10.1038/s41592-019-0654-x

110. McCarthy DJ, Campbell KR, Lun ATL, Wills QF. Scater: pre-processing, quality control, normalization and visualization of single-cell RNA-seq data in R. Bioinformatics. 2017;33(8):1179–1186. doi:10.1093/bioinformatics/btw777

111. Love MI, Huber W, Anders S. Moderated estimation of fold change and dispersion for RNA-seq data with DESeq2. Genome Biology. 2014;15(12):550. doi:10.1186/s13059-014-0550-8

112. L. Lun AT, Bach K, Marioni JC. Pooling across cells to normalize single-cell RNA sequencing data with many zero counts. Genome Biology. 2016;17(1):75. doi:10.1186/s13059-016-0947-7

113. Robinson MD, Oshlack A. A scaling normalization method for differential expression analysis of RNA-seq data. Genome Biology. 2010;11(3):R25. doi:10.1186/gb-2010-11-3-r25

114. Lun ATL, McCarthy DJ, Marioni JC. A step-by-step workflow for low-level analysis of single-cell RNA-seq data with Bioconductor. F1000Res. 2016;5:2122. doi:10.12688/f1000research.9501.2

115. Germain PL, Lun A, Garcia Meixide C, Macnair W, Robinson MD. Doublet identification in single-cell sequencing data using scDblFinder. F1000Res. 2022;10:979. doi:10.12688/f1000research.73600.2

116. Haghverdi L, Lun ATL, Morgan MD, Marioni JC. Batch effects in single-cell RNA-sequencing data are corrected by matching mutual nearest neighbors. Nat Biotechnol. 2018;36(5):421–427. doi:10.1038/nbt.4091

117. Aran D, Looney AP, Liu L, et al. Reference-based analysis of lung single-cell sequencing reveals a transitional profibrotic macrophage. Nat Immunol. 2019;20(2):163–172. doi:10.1038/s41590-018-0276-y

118. Benayoun BA, Pollina EA, Singh PP, et al. Remodeling of epigenome and transcriptome landscapes with aging in mice reveals widespread induction of inflammatory responses. Genome Res. 2019;29(4):697–709. doi:10.1101/gr.240093.118

119. Chen Y, Lun ATL, Smyth GK. From reads to genes to pathways: differential expression analysis of RNA-Seq experiments using Rsubread and the edgeR quasi-likelihood pipeline. F1000Res. 2016;5:1438. doi:10.12688/f1000research.8987.2

120. Müller-Dott S, Tsirvouli E, Vazquez M, et al. Expanding the coverage of regulons from high-confidence prior knowledge for accurate estimation of transcription factor activities. Nucleic Acids Res. 2023;51(20):10934–10949. doi:10.1093/nar/gkad841

121. Badia-i-Mompel P, Vélez Santiago J, Braunger J, et al. decoupleR: ensemble of computational methods to infer biological activities from omics data. Bioinformatics Advances. 2022;2(1):vbac016. doi:10.1093/bioadv/vbac016

122. Ji Z, Ji H. TSCAN: Pseudo-time reconstruction and evaluation in single-cell RNA-seq analysis. Nucleic Acids Research. 2016;44(13):e117. doi:10.1093/nar/gkw430

123. Ahuja G, Antill A, Su Y, et al. Multi-agent AI enables evidence-based cell annotation in single-cell transcriptomics. *bioRxiv*. Preprint posted online November 7, 2025:2025.11.06.686964. doi:10.1101/2025.11.06.686964

124. Ouyang JF, Kamaraj US, Cao EY, Rackham OJL. ShinyCell: simple and sharable visualization of single-cell gene expression data. Bioinformatics. 2021;37(19):3374–3376. doi:10.1093/bioinformatics/btab209

125. Goedhart J, Luijsterburg MS. VolcaNoseR is a web app for creating, exploring, labeling and sharing volcano plots. Sci Rep. 2020;10(1):20560. doi:10.1038/s41598-020-76603-3

126. Ge SX, Jung D, Yao R. ShinyGO: a graphical gene-set enrichment tool for animals and plants. Bioinformatics. 2020;36(8):2628–2629. doi:10.1093/bioinformatics/btz931

127. Landini G, Martinelli G, Piccinini F. Colour deconvolution: stain unmixing in histological imaging. Bioinformatics. 2021;37(10):1485–1487. doi:10.1093/bioinformatics/btaa847

128. Ruifrok AC, Johnston DA. Quantification of histochemical staining by color deconvolution. Anal Quant Cytol Histol. 2001;23(4):291–299.

129. Blake JA, Baldarelli R, Kadin JA, et al. Mouse Genome Database (MGD): Knowledgebase for mouse–human comparative biology. Nucleic Acids Res. 2021;49(D1):D981–D987. doi:10.1093/nar/gkaa1083

