## Supplemental Figures for "Endometrial Hyperplasia Risk Is Increased by High-Fat Diet Via Estrogen-Driven Stromal Fibroblast Reprogramming Toward a Pro-Fibrotic State"

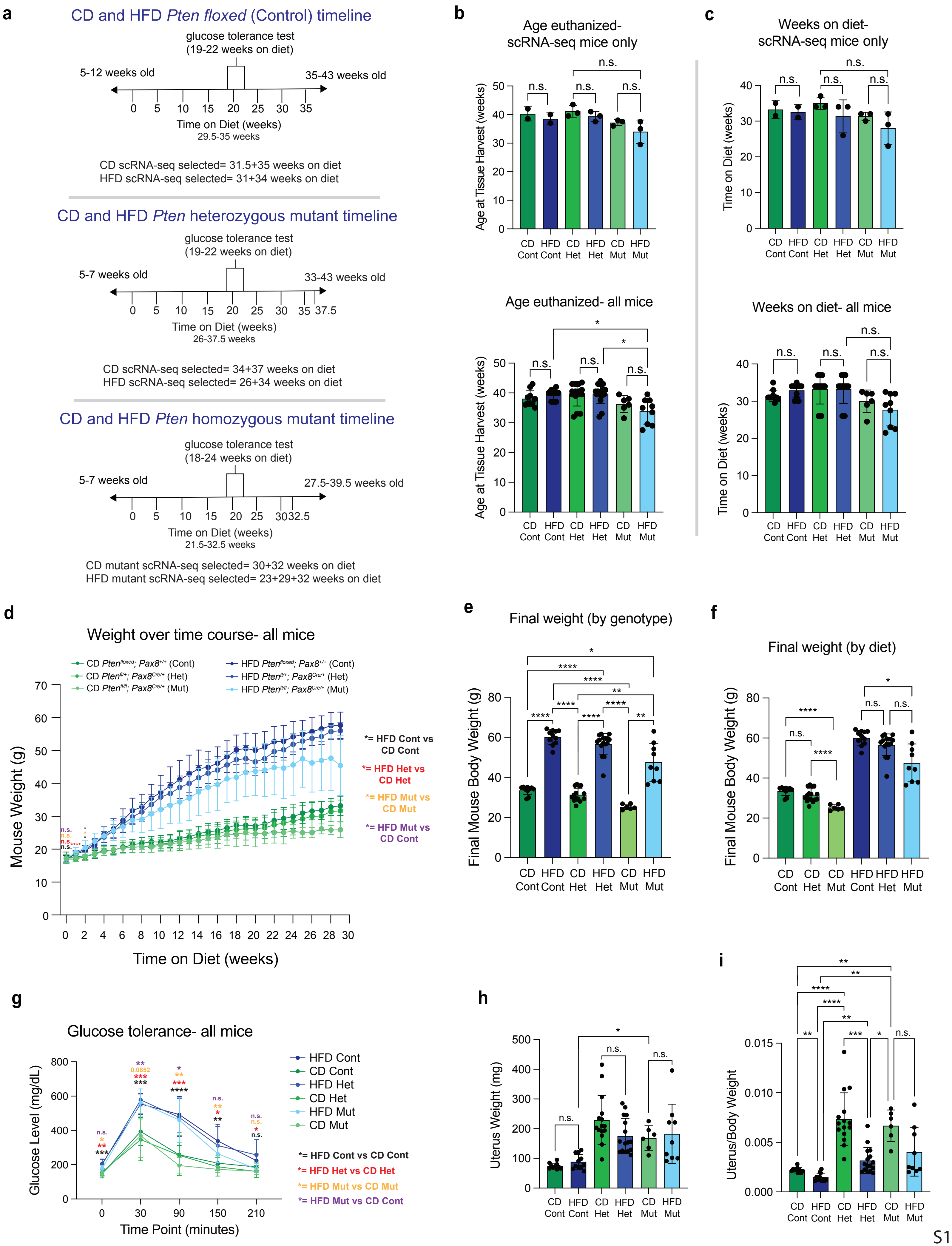
a
CD and HFD Pten floxed (Control) timeline
glucose tolerance test (19-22 weeks on diet)
5-12 weeks old
35-43 weeks old
Time on Diet (weeks)
29.5-35 weeks
CD scRNA-seq selected= 31.5+35 weeks on diet
HFD scRNA-seq selected= 31+34 weeks on diet
CD and HFD Pten heterozygous mutant timeline
glucose tolerance test (19-22 weeks on diet)
5-7 weeks old
33-43 weeks old
Time on Diet (weeks)
26-37.5 weeks
CD scRNA-seq selected= 34+37 weeks on diet
HFD scRNA-seq selected= 26+34 weeks on diet
CD and HFD Pten homozygous mutant timeline
glucose tolerance test (18-24 weeks on diet)
5-7 weeks old
27.5-39.5 weeks old
Time on Diet (weeks)
21.5-32.5 weeks
CD mutant scRNA-seq selected= 30+32 weeks on diet
HFD mutant scRNA-seq selected= 23+29+32 weeks on diet
b
Age euthanized- scRNA-seq mice only
Age euthanized- all mice
Age at Tissue Harvest (weeks)
c
Weeks on diet- scRNA-seq mice only
Weeks on diet- all mice
Time on Diet (weeks)
CD Cont HFD Cont CD Het HFD Het CD Mut HFD Mut
d
Weight over time course- all mice
CD Pten^floxed; Pax8^+/+ (Cont)
CD Pten^fl/+; Pax8^Cre/+ (Het)
CD Pten^fl/fl; Pax8^Cre/+ (Mut)
HFD Pten^floxed; Pax8^+/+ (Cont)
HFD Pten^fl/+; Pax8^Cre/+ (Het)
HFD Pten^fl/fl; Pax8^Cre/+ (Mut)
Mouse Weight (g)
Time on Diet (weeks)
*= HFD Cont vs CD Cont
*= HFD Het vs CD Het
*= HFD Mut vs CD Mut
*= HFD Mut vs CD Cont
e
Final weight (by genotype)
Final Mouse Body Weight (g)
f
Final weight (by diet)
Final Mouse Body Weight (g)
g
Glucose tolerance- all mice
Glucose Level (mg/dL)
Time Point (minutes)
HFD Cont
CD Cont
HFD Het
CD Het
HFD Mut
CD Mut
*= HFD Cont vs CD Cont
*= HFD Het vs CD Het
*= HFD Mut vs CD Mut
*= HFD Mut vs CD Cont
h
Uterus Weight (mg)
i
Uterus/Body Weight

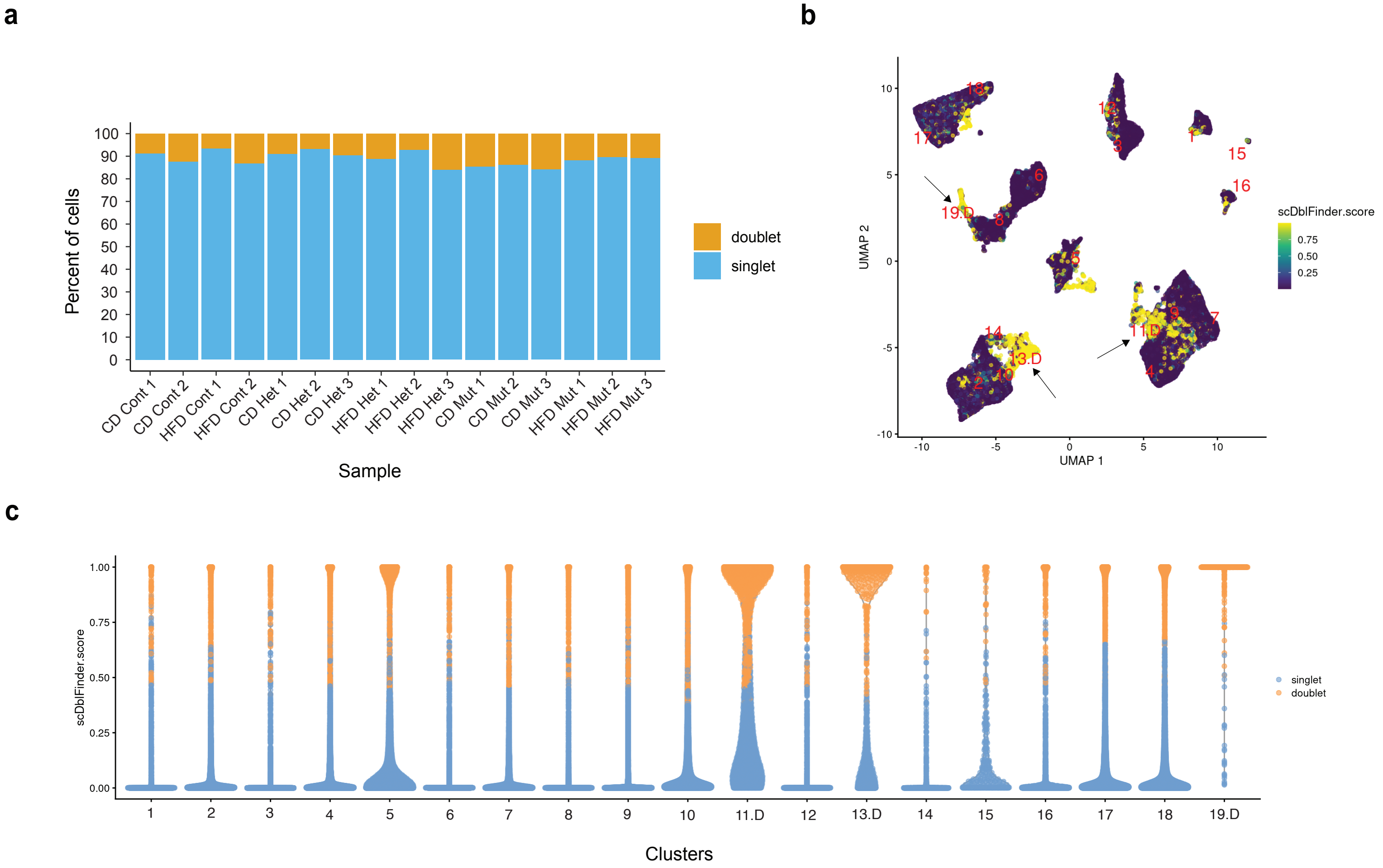

a
b
c
Percent of cells
Sample
doublet
singlet
CD Cont 1
CD Cont 2
HFD Cont 1
HFD Cont 2
CD Het 1
CD Het 2
CD Het 3
HFD Het 1
HFD Het 2
HFD Het 3
CD Mut 1
CD Mut 2
CD Mut 3
HFD Mut 1
HFD Mut 2
HFD Mut 3
UMAP 1
UMAP 2
scDblFinder.score
19.D
11.D
13.D
Clusters

**a**

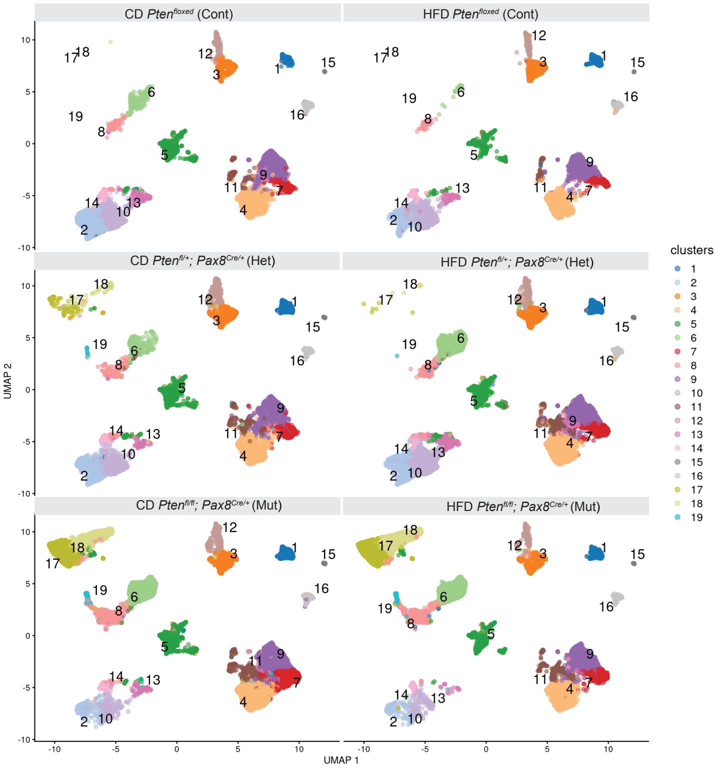


**b**

CD *Pten*$^{fl/fl}$; *Pax8*$^{Cre/+}$ (Mut)
vs *Pten*$^{floxed}$ (Cont)
Cluster abundance*

| Cluster | FC |
|---|---|
| 17 | 12.68 |
| 18 | 9.82 |
| 6 | 3.45 |
| 15 | 2.79 |
| 14 | -1.67 |

HFD *Pten*$^{fl/fl}$; *Pax8*$^{Cre/+}$ (Mut)
vs *Pten*$^{floxed}$ (Cont)
Cluster abundance*

| Cluster | FC |
|---|---|
| 17 | 13.86 |
| 18 | 13.52 |
| 6 | 5.62 |
| 8 | 4.15 |
| 14 | -1.95 |
| 16 | -2.35 |
| 3 | -2.80 |
| 10 | -5.65 |

**c**

### DEGs with diet (in *Pten*$^{floxed}$; *Pax8*$^{Cre/+}$ mice)

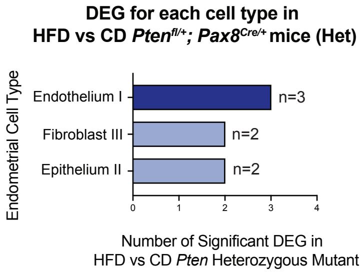


**d**

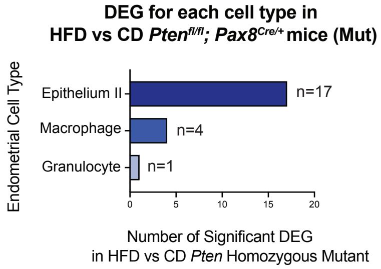


### DEGs with *Pten* mutation (in CD mice)

**e**

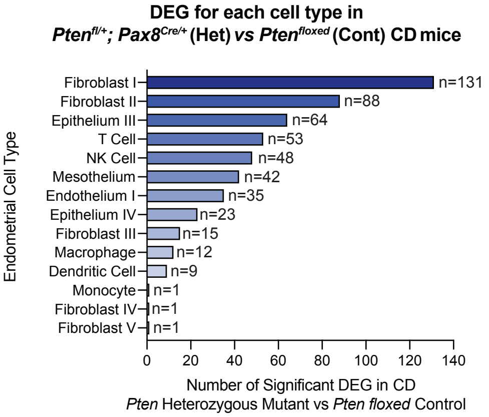


**f**

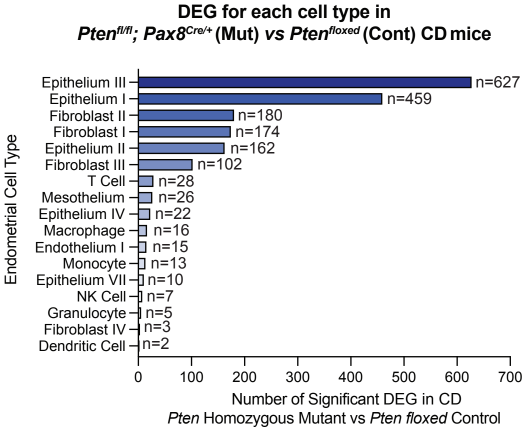


### DEGs with *Pten* mutation (in HFD mice)

**g**

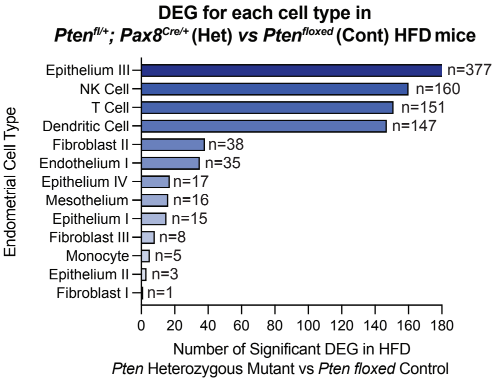


**h**

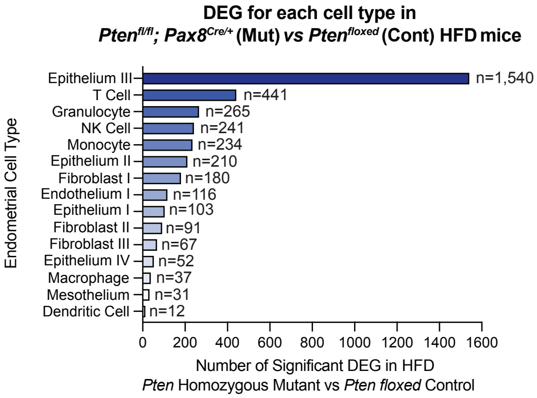

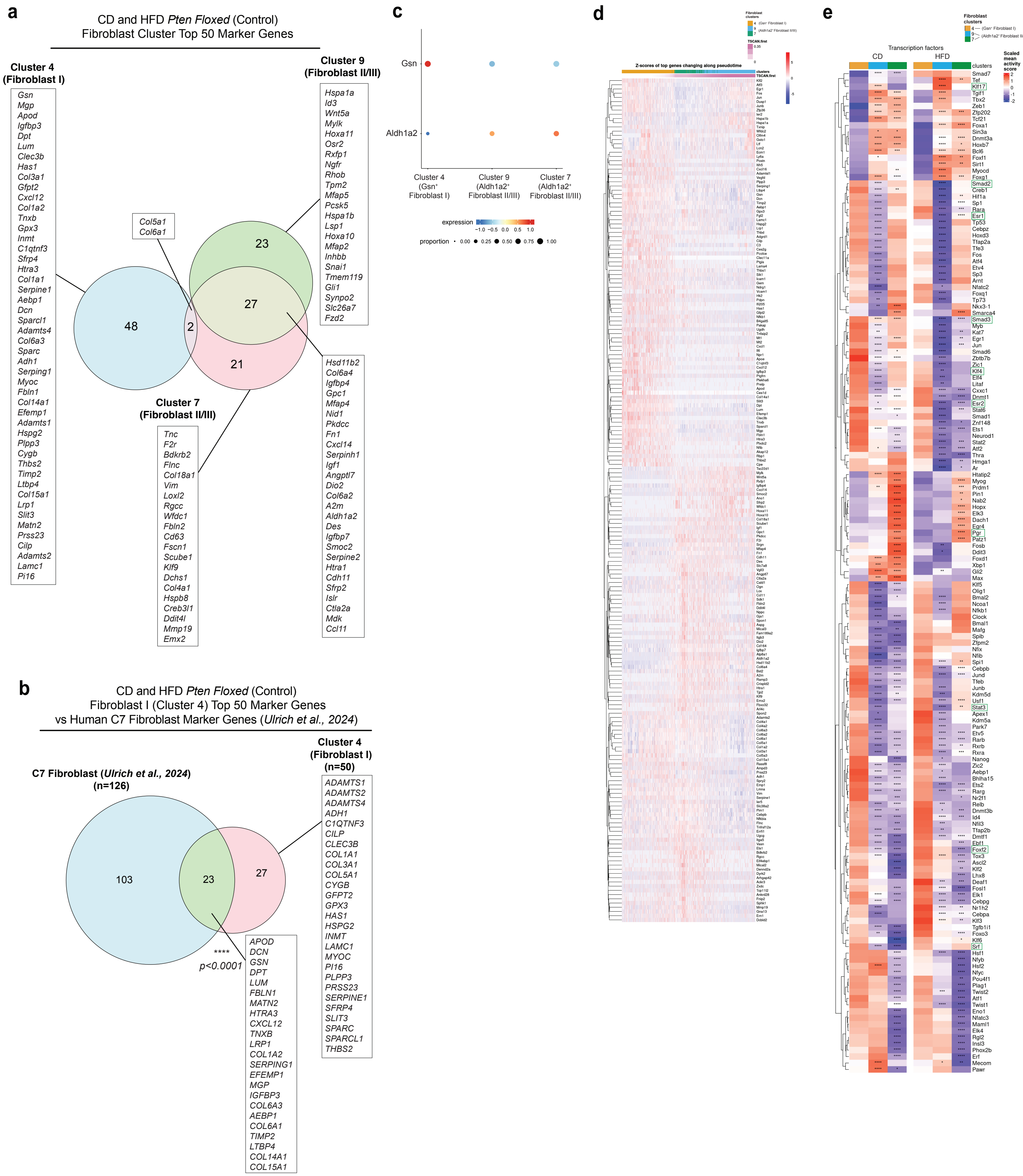
a
CD and HFD Pten Floxed (Control)
Fibroblast Cluster Top 50 Marker Genes
Cluster 4
(Fibroblast I)
Gsn
Mgp
Apod
Igfbp3
Dpt
Lum
Clec3b
Has1
Col3a1
Gfpt2
Cxcl12
Col1a2
Tnxb
Gpx3
Inmt
C1qtnf3
Sfrp4
Htra3
Col1a1
Serpine1
Aebp1
Dcn
Sparcl1
Adamts4
Col6a3
Sparc
Adh1
Serping1
Myoc
Fbln1
Col14a1
Efemp1
Adamts1
Hspg2
Plpp3
Cygb
Thbs2
Timp2
Ltbp4
Col15a1
Lrp1
Slit3
Matn2
Prss23
Cilp
Adamts2
Lamc1
Pi16
Col5a1
Col6a1
48
2
23
27
21
Cluster 9
(Fibroblast II/III)
Hspa1a
Id3
Wnt5a
Mylk
Hoxa11
Osr2
Rxfp1
Ngfr
Rhob
Tpm2
Mfap5
Pcsk5
Hspa1b
Lsp1
Hoxa10
Mfap2
Inhbb
Snai1
Tmem119
Gli1
Synpo2
Slc26a7
Fzd2
Cluster 7
(Fibroblast II/III)
Tnc
F2r
Bdkrb2
Flnc
Col18a1
Vim
Loxl2
Rgcc
Wfdc1
Fbln2
Cd63
Fscn1
Scube1
Klf9
Dchs1
Col4a1
Hspb8
Creb3l1
Ddit4l
Mmp19
Emx2
Hsd11b2
Col6a4
Igfbp4
Gpc1
Mfap4
Nid1
Pkdcc
Fn1
Cxcl14
Serpinh1
Igf1
Angptl7
Dio2
Col6a2
A2m
Aldh1a2
Des
Igfbp7
Smoc2
Serpine2
Htra1
Cdh11
Sfrp2
Islr
Ctla2a
Mdk
Ccl11
b
CD and HFD Pten Floxed (Control)
Fibroblast I (Cluster 4) Top 50 Marker Genes
vs Human C7 Fibroblast Marker Genes (Ulrich et al., 2024)
C7 Fibroblast (Ulrich et al., 2024)
(n=126)
Cluster 4
(Fibroblast I)
(n=50)
103
23
27
****
p<0.0001
APOD
DCN
GSN
DPT
LUM
FBLN1
MATN2
HTRA3
CXCL12
TNXB
LRP1
COL1A2
SERPING1
EFEMP1
MGP
IGFBP3
COL6A3
AEBP1
COL6A1
TIMP2
LTBP4
COL14A1
COL15A1
ADAMTS1
ADAMTS2
ADAMTS4
ADH1
C1QTNF3
CILP
CLEC3B
COL1A1
COL3A1
COL5A1
CYGB
GFPT2
GPX3
HAS1
HSPG2
INMT
LAMC1
MYOC
PI16
PLPP3
PRSS23
SERPINE1
SFRP4
SLIT3
SPARC
SPARCL1
THBS2
c
Gsn
Aldh1a2
Cluster 4 (Gsn⁺ Fibroblast I)
Cluster 9 (Aldh1a2⁺ Fibroblast II/III)
Cluster 7 (Aldh1a2⁺ Fibroblast II/III)
expression
−1.0 −0.5 0.0 0.5 1.0
proportion
0.00 0.25 0.50 0.75 1.00
d
Z-scores of top genes changing along pseudotime
clusters
TSCAN.first
e
Transcription factors
CD
HFD
clusters
Scaled mean activity score

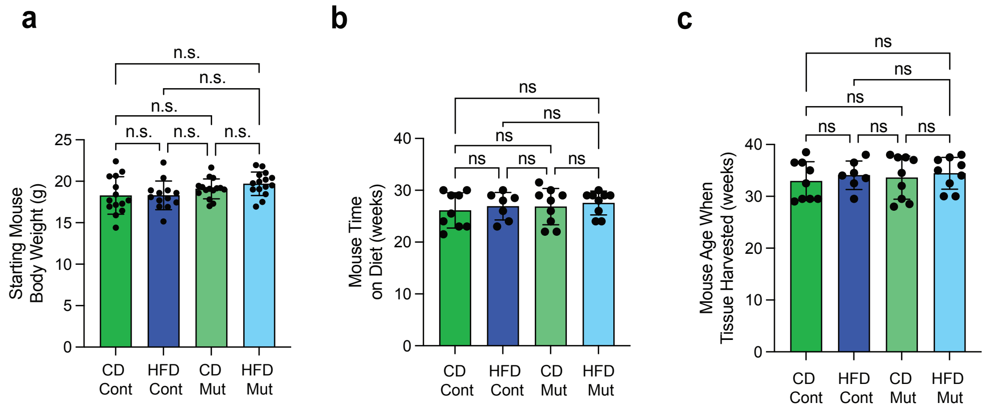
a
b
c
n.s.
ns
Starting Mouse Body Weight (g)
Mouse Time on Diet (weeks)
Mouse Age When Tissue Harvested (weeks)
0
5
10
15
20
25
30
40
CD Cont
HFD Cont
CD Mut
HFD Mut

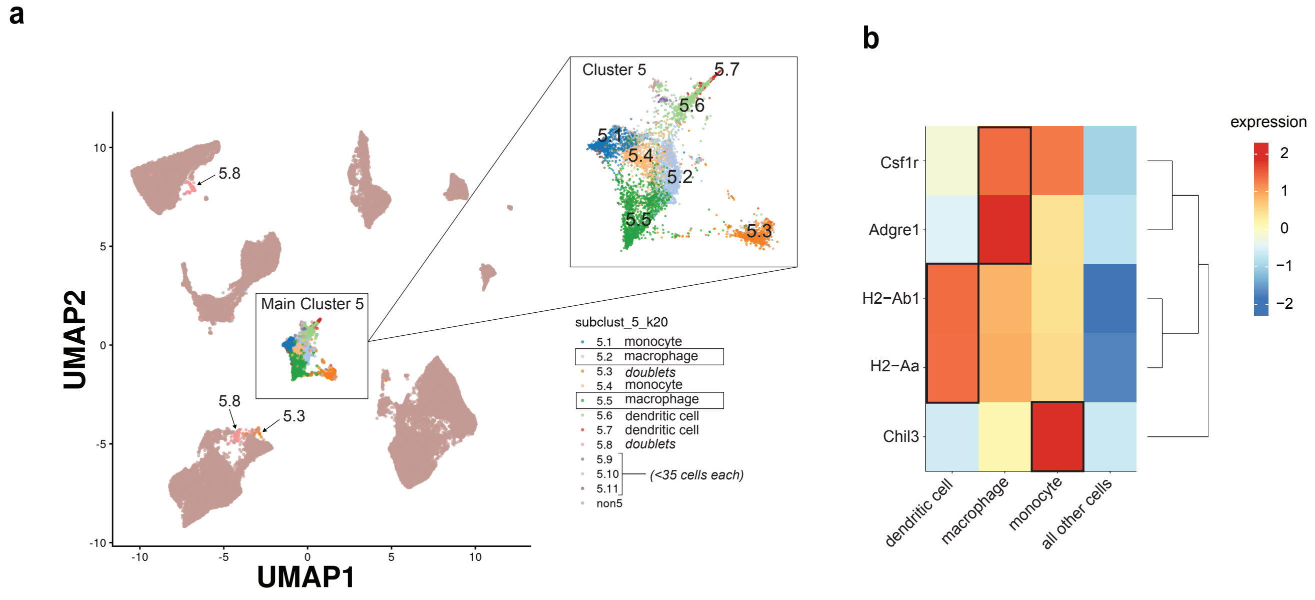
a
b
Cluster 5
5.7
5.6
5.1
5.4
5.2
5.5
5.3
5.8
Main Cluster 5
UMAP2
UMAP1
subclust_5_k20
5.1 monocyte
5.2 macrophage
5.3 doublets
5.4 monocyte
5.5 macrophage
5.6 dendritic cell
5.7 dendritic cell
5.8 doublets
5.9
5.10
5.11
(<35 cells each)
non5
expression
Csf1r
Adgre1
H2−Ab1
H2−Aa
Chil3
dendritic cell
macrophage
monocyte
all other cells

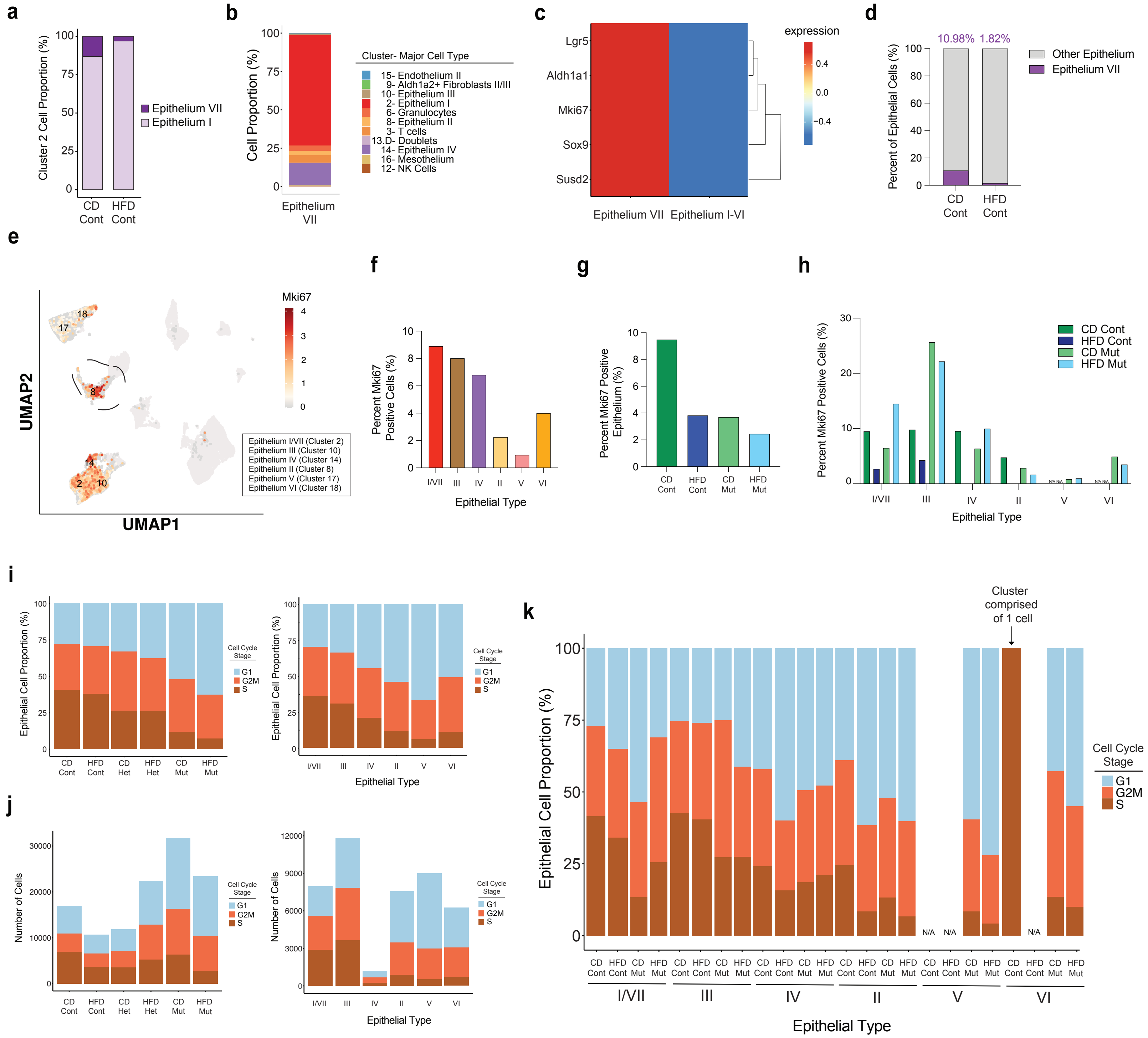

a
Cluster 2 Cell Proportion (%)
Epithelium VII
Epithelium I
CD Cont
HFD Cont
b
Cell Proportion (%)
Cluster- Major Cell Type
15- Endothelium II
9- Aldh1a2+ Fibroblasts II/III
10- Epithelium III
2- Epithelium I
6- Granulocytes
8- Epithelium II
3- T cells
13.D- Doublets
14- Epithelium IV
16- Mesothelium
12- NK Cells
Epithelium VII
c
expression
Lgr5
Aldh1a1
Mki67
Sox9
Susd2
Epithelium VII
Epithelium I-VI
d
10.98%
1.82%
Percent of Epithelial Cells (%)
Other Epithelium
Epithelium VII
CD Cont
HFD Cont
e
Mki67
UMAP2
UMAP1
Epithelium I/VII (Cluster 2)
Epithelium III (Cluster 10)
Epithelium IV (Cluster 14)
Epithelium II (Cluster 8)
Epithelium V (Cluster 17)
Epithelium VI (Cluster 18)
f
Percent Mki67 Positive Cells (%)
Epithelial Type
g
Percent Mki67 Positive Epithelium (%)
CD Cont
HFD Cont
CD Mut
HFD Mut
h
Percent Mki67 Positive Cells (%)
CD Cont
HFD Cont
CD Mut
HFD Mut
Epithelial Type
i
Epithelial Cell Proportion (%)
Cell Cycle Stage
G1
G2M
S
CD Cont
HFD Cont
CD Het
HFD Het
CD Mut
HFD Mut
Epithelial Type
j
Number of Cells
Cell Cycle Stage
Epithelial Type
k
Cluster comprised of 1 cell
Epithelial Cell Proportion (%)
Cell Cycle Stage
N/A
Epithelial Type

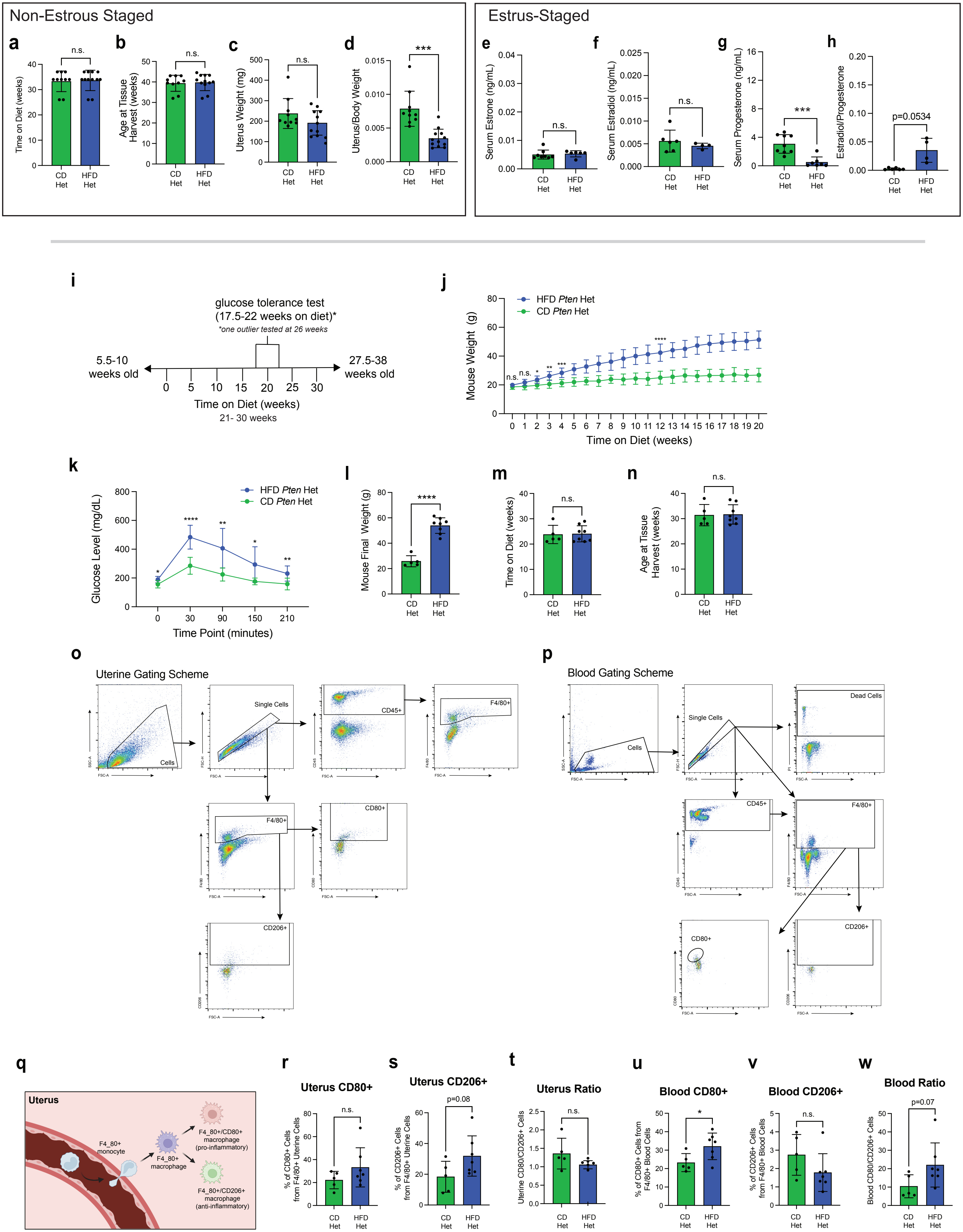
Non-Estrous Staged
Estrus-Staged
a
b
c
d
e
f
g
h
Time on Diet (weeks)
Age at Tissue Harvest (weeks)
Uterus Weight (mg)
Uterus/Body Weight
Serum Estrone (ng/mL)
Serum Estradiol (ng/mL)
Serum Progesterone (ng/mL)
Estradiol/Progesterone
n.s.
***
p=0.0534
CD Het
HFD Het
i
glucose tolerance test
(17.5-22 weeks on diet)*
*one outlier tested at 26 weeks
5.5-10 weeks old
27.5-38 weeks old
Time on Diet (weeks)
21- 30 weeks
j
HFD Pten Het
CD Pten Het
Mouse Weight (g)
Time on Diet (weeks)
k
Glucose Level (mg/dL)
Time Point (minutes)
l
Mouse Final Weight (g)
m
Time on Diet (weeks)
n
Age at Tissue Harvest (weeks)
o
Uterine Gating Scheme
Cells
Single Cells
CD45+
F4/80+
CD80+
CD206+
p
Blood Gating Scheme
Dead Cells
q
Uterus
F4_80+ monocyte
F4_80+ macrophage
F4_80+/CD80+ macrophage (pro-inflammatory)
F4_80+/CD206+ macrophage (anti-inflammatory)
r
Uterus CD80+
% of CD80+ Cells from F4/80+ Uterine Cells
s
Uterus CD206+
% of CD206+ Cells from F4/80+ Uterine Cells
p=0.08
t
Uterus Ratio
Uterine CD80/CD206+ Cells
u
Blood CD80+
% of CD80+ Cells from F4/80+ Blood Cells
v
Blood CD206+
% of CD206+ Cells from F4/80+ Blood Cells
w
Blood Ratio
Blood CD80/CD206+ Cells
p=0.07

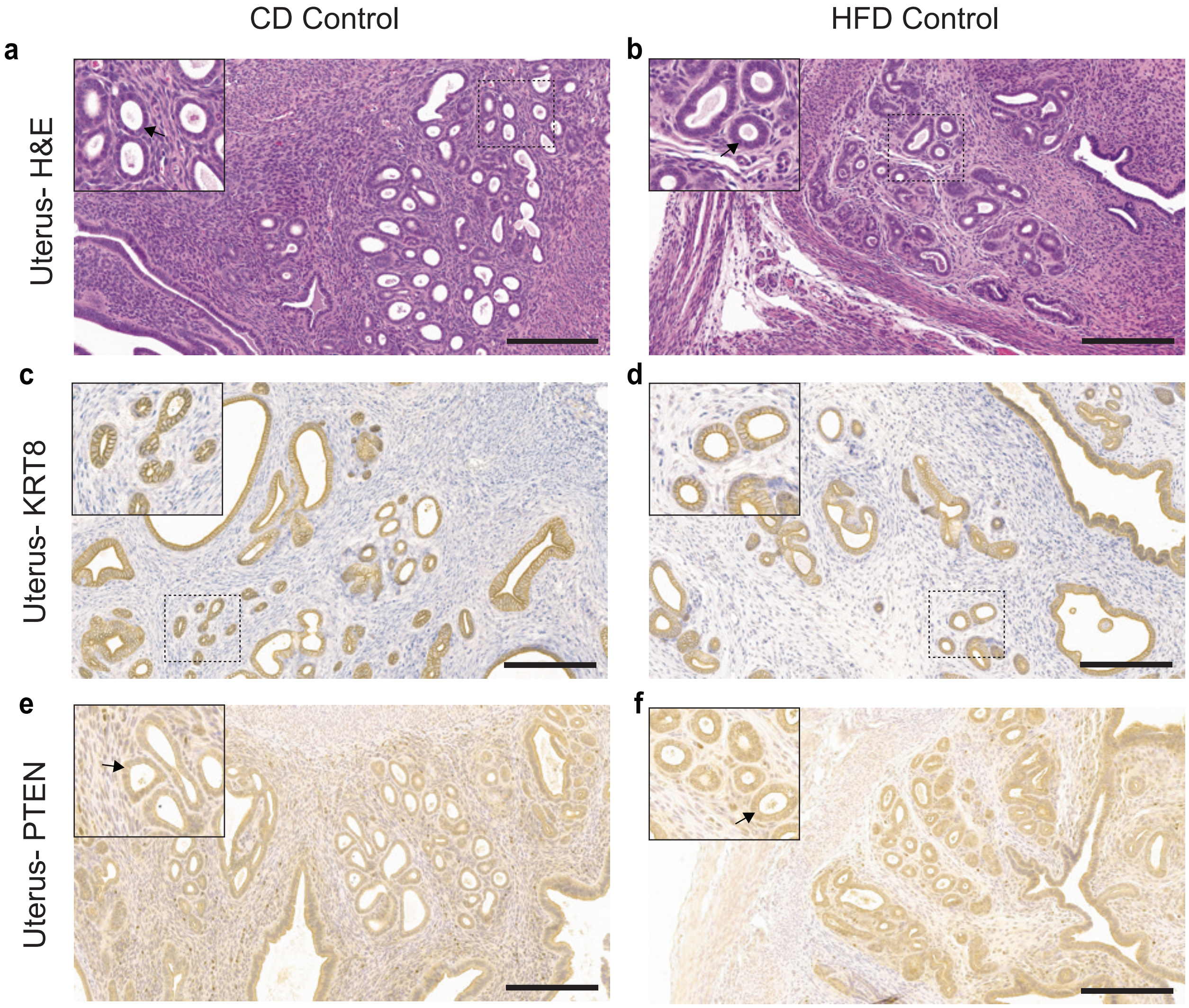

CD Control
HFD Control
a
b
Uterus- H&E
c
d
Uterus- KRT8
e
f
Uterus- PTEN
